# Identification of a distinct ductal subpopulation with self-renewal and differentiation potential from the adult murine pancreas

**DOI:** 10.1101/2022.07.12.499653

**Authors:** Jacob R. Tremblay, Jose Ortiz, Janine C. Quijano, Heather Zook, Jeanne M. LeBon, Wendong Li, Kevin Jou, Walter Tsark, Jeffrey R. Mann, Mark Kozlowski, David A. Tirrell, Farzad Esni, Dannielle D. Engle, Arthur D. Riggs, Hsun Teresa Ku

## Abstract

Pancreatic ducts function to deliver digestive enzymes into the intestines. Upon injury, ducts can become proliferative and contribute to tissue regeneration; however, the identity of the ductal cells that contribute to these processes is unknown. We combined fluorescence-activated cell sorting, a methylcellulose-containing 3-dimensional culture, droplet RNA-sequencing, and a clonal lineage tracing tool to identify and isolate a distinct subpopulation of pancreatic ductal cells that exhibit progenitor cell properties. These ductal cells are unique in that they form tightly-bound clusters (termed FSC^mid-high^), with an average of 8 cells per cluster. FSC^mid-high^ clusters comprise only about 0.1% of the total pancreas, are tri-potent for duct, acinar and endocrine lineages, and self-renew robustly *in vitro*. Transcriptomic analysis of FSC^mid-high^ clusters reveals enrichment for genes involved in cell-cell interactions, organ development, and cancer pathways. FSC^mid-high^ clusters express embryonic pancreatic progenitor markers Sox9, Pdx1, and Nkx6-1 at both transcription and protein levels. FSC^mid-high^ clusters are resistant to enzymatic dissociation and survive severe *in vivo* acinar injury, which induces formation of ductal rosettes that become proliferative within 14 days. Thus, FSC^mid-high^ clusters represent a small subset of ductal cells with progenitor cell properties. These rare progenitor-like duct cell clusters have implications in pancreas regeneration and tumor initiation/progression.

## Introduction

In the adult pancreatic epithelium, there are three major lineages of cells: acinar, duct, and endocrine cells. Acinar and ductal cells are responsible for secreting and transporting digestive enzymes, respectively, to aid in nutrient digestion while endocrine cells secrete hormones to regulate glucose homeostasis. Adult pancreatic cells are mostly quiescent during steady state ^1,2^. However, when damage and stress occur to acinar and endocrine insulin-producing beta cells ^3–6^, which results in pancreatitis and diabetes, respectively, proliferation increases in not only acinar and endocrine cells but also ductal cells ^7,8^. Ductal cells have been implicated as a source of progenitor cells that contribute to beta cell neogenesis ^9,10^ or pancreatic adenocarcinoma ^11^. Although the roles of adult ductal cells as progenitor cells are still controversial ^12–15^, recent emerging evidence has strongly suggested that certain subpopulation of ductal cells are involved in regeneration of endocrine cells in conditions of insulin resistance ^16^ and insulin-dependent diabetes ^17,18^. However, the fundamental identifies of these progenitor-like ductal cells remain unknown.

To identify potentially rare adult ductal progenitor cells, here we employed an unbiased fractionation strategy by using fluorescence-activated cell sorting (FACS) on dissociated murine pancreatic cells; we discover a tightly-bound ductal cell cluster (termed FSC^mid-high^ cluster), which constitutes only 0.1% of the total pancreatic cells, as the fundamental progenitor-like cells in the normal adult murine pancreas. The FSC^mid-high^ clusters can self-renew and differentiate *in vitro* and are resistant to *in vivo* acinar injury conditions. Our results highlight the highly heterogeneous nature of the adult ductal cells, which may explain the difficulties in studying these rare ductal cells in the past.

## Results

### CD133^high^CD71^low^ duct cells contain cell fractions with distinct morphologies

Flow cytometry can be used to distinguish individual cells or small clusters based on light scattering at different angles: the detected light beam in line with the originating beam is defined as forward scatter (FSC), and that in perpendicular as side scatter (SSC). FSC is indicative of particle size while SSC complexity or granularity, and the combination of these two parameters has been useful to distinguish various cell types ^19^. We hypothesized that adult pancreatic progenitor cells may be small in size with a high nuclear-to-cytoplasm ratio, similar to adult hematopoietic stem cells ^20,21^, owing to a generally quiescent state in homeostasis ^22^. To test this hypothesis, whole pancreata from adult mice were dissociated and analyzed by flow cytometry.

We previously established that a ductal cell sub-population identified by CD133^high^CD71^low^ cell-surface staining shows progenitor-like properties *in vitro*^23^. We therefore further analyzed the CD133^high^CD71^low^ duct population based on FSC and SSC (Fig. 1a and Supplementary Fig. 1). Four sub-populations emerged from the parent CD133^high^CD71^low^ population: abbreviated hereafter as FSC^low^, FSC^mid-low^, FSC^mid-high^, and FSC^high^. These 4 sub-populations constituted 1.27 ± 0.27, 0.47 ± 0.13, 0.12 ± 0.04 and 0.23 ± 0.16% of the total pancreatic population, and 53.5 ± 7.8, 28.2 ± 2.4, 5.3 ± 0.5 and 7.5 ± 5.8% of the gated CD133^high^CD71^low^ population, respectively. Adding the % total pancreatic cells from all 4 sub-populations amounted to 2.09%, which is within the range of the parent CD133^high^CD71^low^ population among total pancreatic cells reported in our prior study (average 2.4 ± 1.9%) ^23^. To examine cell morphology, freshly-sorted sub-populations were cytospun and stained with Wright-Giemsa to distinguish nuclei and cytoplasm. The FSC^mid-high^ sub-population contained aggregates of cells with high nuclear-to-cytoplasm ratios that were highly resistant to enzymatic single cell dissociation (to be addressed later in the study), which we named “small clusters” (Fig 1b; upper right panel). The FSC^low^ sub-population was comprised of mostly single cells with high nuclear-to-cytoplasm ratios, which we named “single” (Fig. 1b; upper left panel). The “large” morphology in the FSC^high^ population had two phenotypes: one was bi-nucleated with pink cytoplasm (Fig. 1b; lower right panel), suggestive of some acinar cells ^24^ expressing CD133^high^ on the cell surface, and the other was single-nucleated with purple cytoplasm. The identity of the latter cell is currently unknown. The FSC^mid-low^ population contained a mixture of the aforementioned 3 cell morphologies, with mostly “elongated” morphology plus small clusters (Fig. 1b; lower left panel). These 4 morphologies were also identifiable under a phase-contrast light microscope (Fig. 1c). The diameters of these morphologies were measured; “single” had the smallest diameters, while “elongated”, “small clusters”, and “large” had increasingly larger diameters (Fig. 1d). Counting the 4 morphologies in each sorted sub-population revealed that higher FSC was positively correlated with an increased proportion of morphologies with larger diameters (Fig. 1e), confirming effective sorting based on FSC.

**Figure 1.**
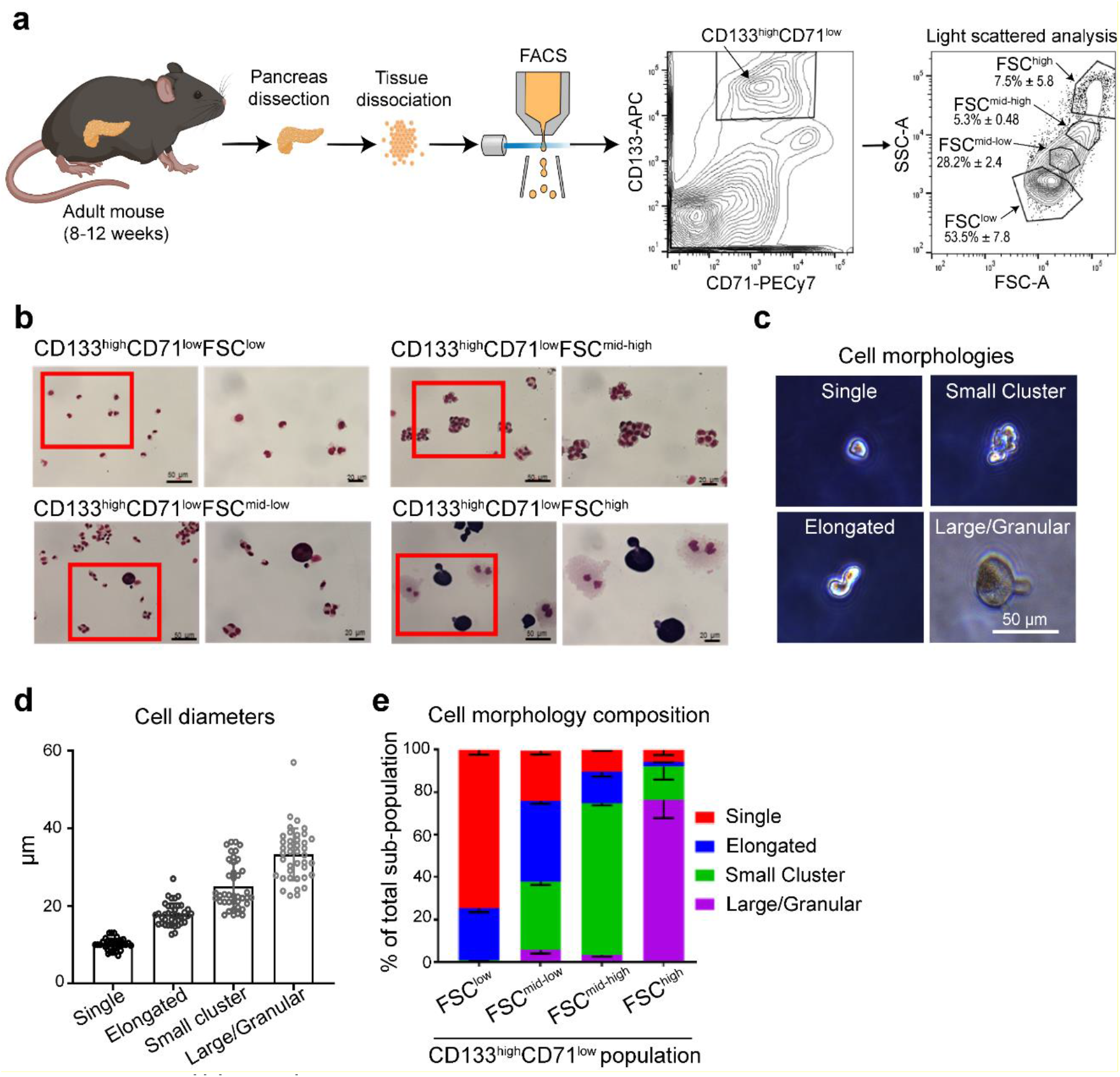
CD133^high^CD71^low^ ductal population consists of forward scatter (FSC) fractions with distinguishable cell morphology. (a) Schematic of experimental design with representative flow cytometry analysis. (b) Representative photomicrographs of sorted sub-populations that were stained with modified Wright-Giemsa solution. Scale bars on the lower and higher magnification represent 50 and 20 μm, respectively. (c) Representative photomicrographs of freshly-sorted four cell morphologies detected in the parent CD133^high^CD71^low^ population observed under a phase-contrast light microscope. Scale bar=50 μm. (d) Measurement of diameters revealed that different morphologies had sizes consistent with their forward scatter in flow cytometry analysis. n=2 independent experiments from 5 mice each and at least 20 replicates in each group. Error bars represent SD. (e) Frequencies of each morphology in each sorted sub-population. Small clusters are most enriched in the CD133^high^CD71^low^FSC^mid-high^ fraction (abbreviated as FSC^mid-high^). n=3 independent sorted experiments for each gate, with 3 different fields of view when counting cell types. Error bars represent SEM.

### The FSC^mid-high^ cell fraction is enriched for pancreas colony-forming units (PCFUs)

The 3D colony assay developed in our laboratory utilizes methylcellulose as a matrix ^25^. Methylcellulose is a biologically inert material ^26,27^ that prevents reaggregation of the plated cells while allowing for expansion into a colony. The advantage of using methylcellulose is that other basement membrane extracts, such as Matrigel, or defined ECM proteins, such as laminin, can be added at lower percentages below their gelling point ^25^. We named a progenitor cell capable of giving rise to a colony a “pancreatic colony-forming unit (PCFU)”, following benchmarks used by hematologists to quantify hematopoietic stem cell activity according to their colony-forming efficiency ^28^.

To quantify the PCFUs in each sub-population, freshly-sorted cells from each fraction were plated into a colony assay (Fig. 2a). In the colony assay containing Matrigel (5% vol/vol) and Rspondin-1 (RSPO1) ^25^ (herein Matrigel/RSPO1 colony assay), the FSC^mid-high^ fraction had the highest PCFUs, as seen by the resulting colonies (total number of 3-week-old colonies, which we called “Cystic”, among 500 plated units) (Fig. 2b). This result contrasted our original hypothesis that the FSC^low^ fraction was most consistent with a stem cell morphology. The FSC^low^ fraction contained some PCFUs, but the colony-forming efficiency was approximately 7-fold lower than the FSC^mid-high^ population (Fig. 2b).

**Figure 2.**
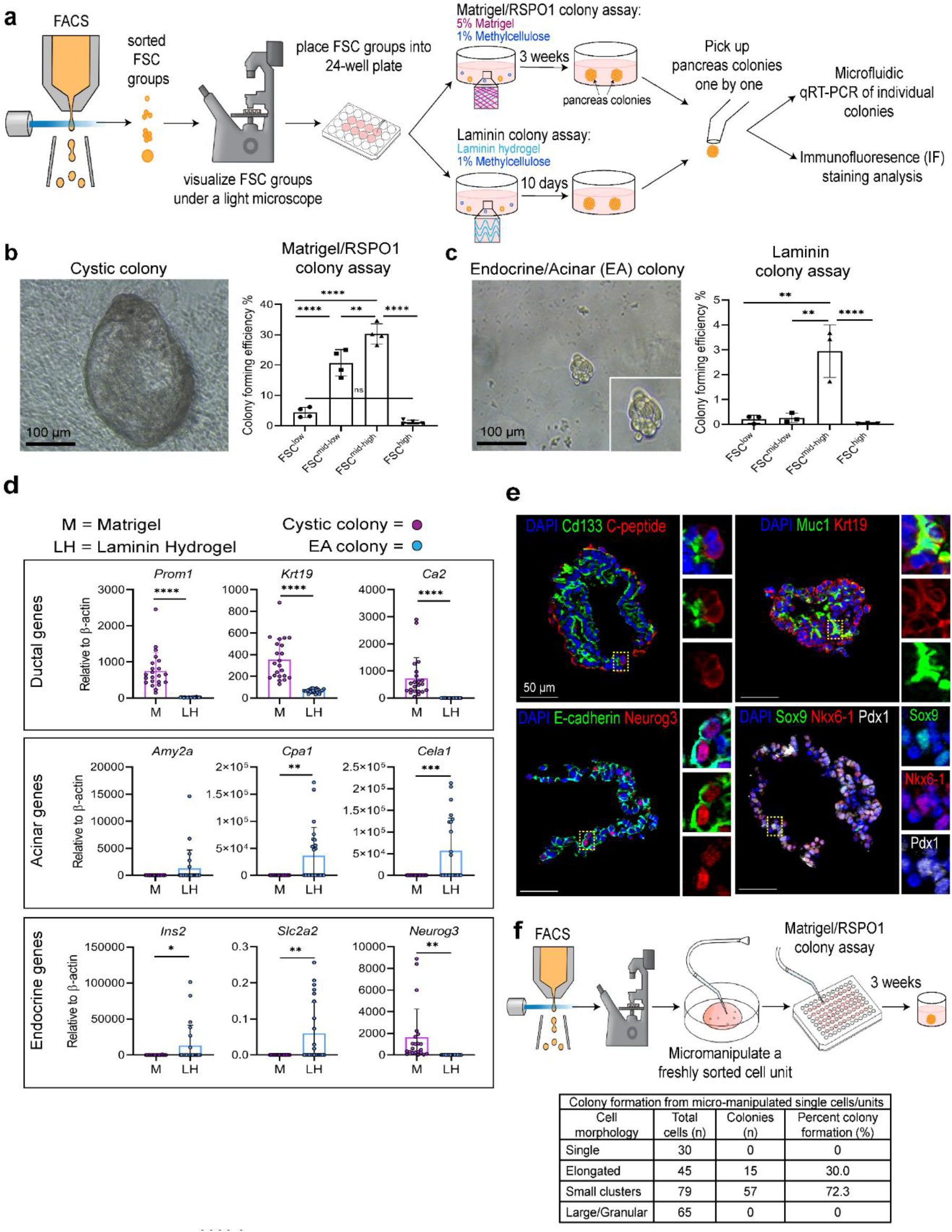
The FSC^mid-high^ fraction is enriched for tri-potent, colony-forming units. (a) Schematic of experimental design with sorted units placed into either the Matrigel/RSPO1 colony assay (which contains 5% Matrigel with 1% Methylcellulose) or the Laminin colony assay (which contains laminin hydrogel with 1% Methylcellulose) followed by culture time to produce colonies for subsequent analysis. (b) Representative image of a Cystic colony grown in Matrigel/RSPO1 colony assay (left). Colony-forming efficiency was determined for each FSC fraction. **p<0.01, ****p<0.0001, n=5. (c) Representative image of an Endocrine/Acinar (EA) colony grown in laminin colony assay (left). Colony-forming efficiency was determined for each FSC fraction. **p<0.01, ****p<0.0001, n=3. Statistics were performed using one-way ANOVA multiple comparisons two-tailed Student’s *t*-test. Error bars represent SEM. (d) Microfluidic qRT-PCR analysis show that individual FSC^mid-high^-derived colonies grown in Matrigel (M) and laminin hydrogel (LH) expressed tri-lineage markers. Matrigel/RSPO1 colony assay promotes expression of ductal markers while laminin assay promotes expression of acinar and endocrine markers, consistent with our prior findings ^25,29^. This result demonstrates the tri-lineage differentiation ability of the sorted FSC^mid-high^ sub-population. Each dot represents a colony. *p<0.05, **p<0.01, ***p<0.001, ****p<0.0001, n=22-23. (e) Immunofluorescent (IF) staining of FSC^mid-high^-derived colonies grown in Matrigel/RSPO1 colony assay. Co-staining of epithelial (E-cadherin), ductal (Cd133, Muc1, Krt19), endocrine (C-peptide, Neurog3), and pancreatic progenitor (Sox9, Nkx6-1, Pdx1) markers. Yellow dash boxes are zoomed to the right side. Scale bar= 50 μm. (f) Schematic of single-unit micro-manipulation of sorted morphologies (Single, Elongated, Small clusters, Large/Granular) plated in Matrigel/RSPO1 colony assay. Among micro-manipulated Small clusters and Elongated, 72.3% and 30% gave rise to pancreas colonies, respectively.

We previously showed that performing the colony assay using a hydrogel containing the IKVAV sequence from laminin (herein laminin colony assay) in the absence of Matrigel and RSPO1 led to increased endocrine and acinar lineage markers and decreased ductal lineage markers ^25,29^. These colonies were termed endocrine/acinar (E/A) colonies. In the laminin colony assay, the FSC^mid-high^ fraction also had the greatest number of PCFUs (Fig. 2c), with an approximately 250-fold higher colony-forming efficiency than the FSC^low^ fraction. Because the FSC^mid-high^ fraction has the highest colony forming efficiency, we focused our attention on the FSC^mid-high^ fraction for subsequent analyses.

To determine lineage marker expression, we micro-manipulated and picked one-by-one individual Cystic and E/A colonies and analyzed gene expression by microfluidic qRT-PCR analysis. Consistent with our prior findings ^25,29^, FSC^mid-high^ derived colonies grown in the Matrigel/RSPO1 relative to laminin colony assay expressed higher levels of ductal (*Prom1, Krt19, Ca2*) and endocrine progenitor (*Neurog3*) markers, and lower levels of acinar (*Cpa1, Cela1*) and endocrine cell (*Ins2, Glut2*) markers (Fig. 2d). These results suggest that the FSC^mid-high^ fraction contains multipotent progenitors or lineage-restricted progenitor cells. Protein expression of CD133, C-peptide, Muc1, Krt19, E-cadherin, and Neurog3 was confirmed in the colonies (Fig. 2e and Supplementary Fig. 2). Previous studies determined that co-expression of Sox9, Nkx6-1, Pdx1 in embryonic pancreatic multipotent progenitor cells (MPCs) is necessary for pancreas development ^30,31^. We found cells in the colonies simultaneously expressing these three proteins (Fig. 2e; lower right panel), mimicking pancreatic MPCs during organogenesis.

To determine which cell morphology was responsible for forming colonies, we micro-manipulated FSC populations into Matrigel/RSPO1 colony assay at 1 cell (or cluster) per well in 96-well plates (Fig. 2f). The small round and large granular cells did not form colonies, whereas elongated cells had a 30% colony-forming efficiency and small clusters a notable 72.3% efficiency. Altogether, these data suggest that the multicellular clusters in the FSC^mid-high^ fraction contain functional progenitor cells.

### PCFUs within the FSC^mid-high^ fraction self-renew

One of the defining properties of a progenitor cell is its ability to self-renew. To test the self-renewal potential of the PCFU-enriched FSC^mid-high^ fraction, a serial dissociation and replating strategy was used (Fig. 3a). In our previous studies, we optimized self-renewal conditions in our Matrigel/RSPO1 colony assay, in which exogenous RSPO1 is necessary for PCFU self-renewal ^25^. The FSC^mid-high^ fraction was plated into the Matrigel/RSPO1 colony assay. The resulting 3-week-old primary colonies were dissociated and serially replated for a total of 4 generations. In parallel experiments, we used the FSC^low^ fraction as a control due to its higher proportion among parent cells (Fig. 1a) and lower PCFUs (Fig. 2b-c). The FSC^mid-high^ fraction grew exponentially from the first generation and gave rise to a higher number of total cells (both PCFUs and non-PCFUs) across all generations compared to the FSC^low^ fraction (Fig. 3b-c). Although the FSC^low^ fraction had a lag phase in the early passages, the growth rate (slope of the curve) caught up with the FSC^mid-high^ fraction in the later passages (Fig. 3b-c). This “catch up” effect in later passages of the FSC^low^ population was in both the proportion of the total cells (Fig. 3d) and total colonies (Fig. 3e) per well. Thus, even though the FSC^low^ fraction had fewer PCFUs to begin with, which could be elongated and/or small clusters, the PCFUs retain the same self-renewal capacity as those in the FSC^mid-high^. These results are consistent with the FSC^mid-high^ fraction being more enriched in PCFUs compared to the FSC^low^ fraction (Fig. 2b-c). Overall, after 9 weeks in the self-renewing culture condition, the total cell number and total PCFUs contained in the FSC^mid-high^ fraction expanded ~440,000 and ~78,000 fold, respectively (Fig. 3b-c). This fold expansion of PCFUs is comparable to the sorted CD133^+^Sox9^+^ total ductal cells from our earlier report ^25^.

**Figure 3.**
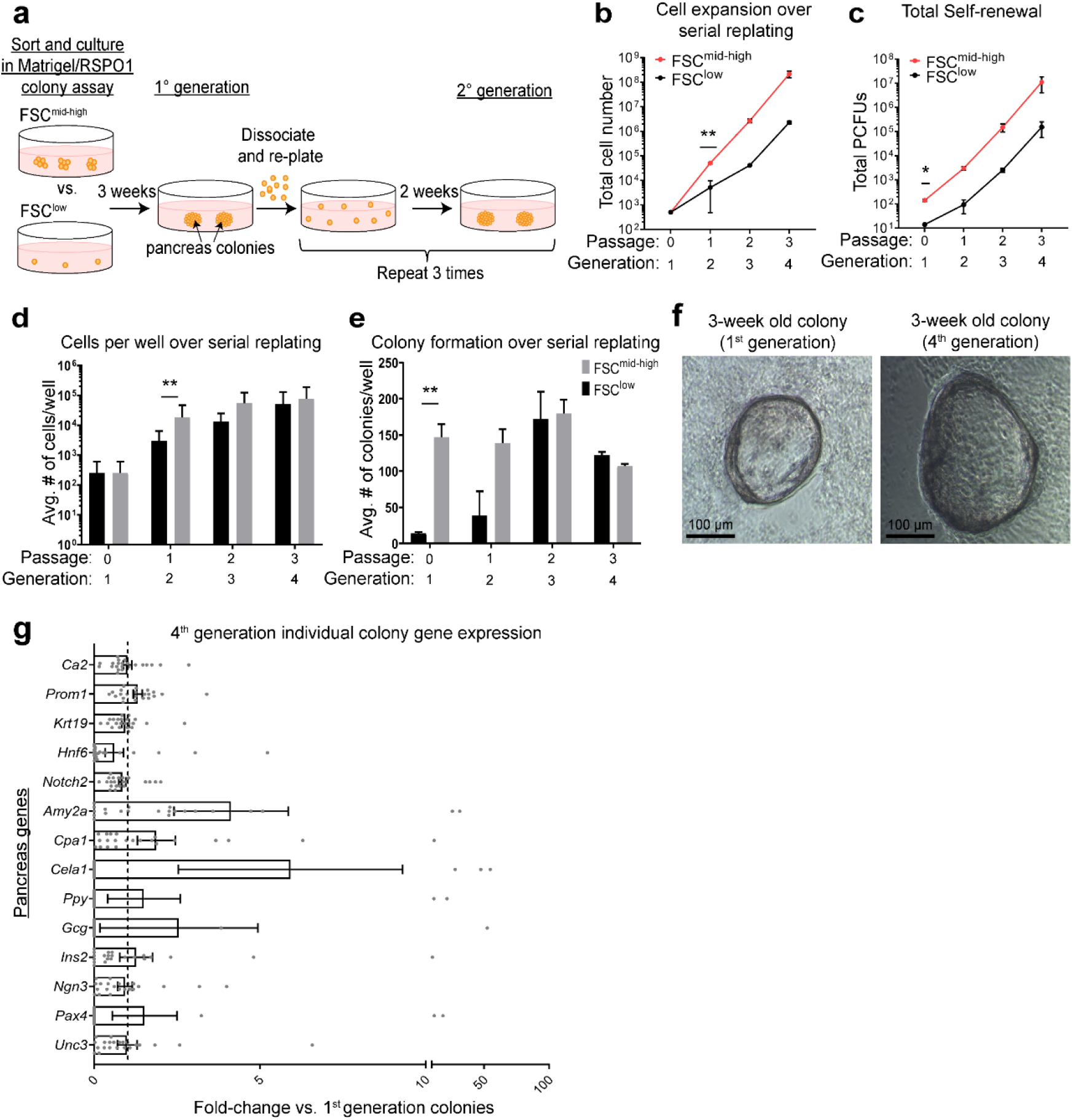
FSC^mid-high^-derived colonies exhibit self-renewal capacity. (a) Schematic of serial replating strategy to assess self-renewal capacity of FSC^mid-high^- or FSC^low^-derived pancreas colonies using the Matrigel/RSPO1 colony assay. Three-week-old colonies grown in the primary (1°) culture were pooled and dissociated. A portion was re-plated into a secondary (2°) culture to generate colonies in 2 weeks. This dissociation and replating were repeated for a total of 3 times. (b-c) Growth curves reveal that pancreatic colony-forming units (PCFUs) from either FSC^low^ or FSC^mid-high^ fractions can self-renew over time, shown by increased overall numbers of total cells (b) and total PCFUs (c). *p<0.05, **p<0.01, n=2 independent experiments consisting of 3 independent replicates each. Error bars represent SEM. (d) Average number of cells per well over serial replating. (e) Average number of colonies per well over serial replating. (f) Photomicrographs of representative 3-week-old Cystic colonies from the 1^st^ and the 4^th^ generations. (g) Microfluidic qRT-PCR analysis of ductal (*CAII, CD133, CK19*) (*Amylase, CPA1, Elastase1*), endocrine (*PPY, Gcg, Ins2, Ngn3*), and other pancreas markers (*Hnf6, Notch2, Pax4, Pdx1, Nkx6-1, Sox9, Unc3*) on individual colonies. Data represent fold-change calculation of gene expression of the 4^th^-generation compared to the 1^st^-generation. **p<0.01, n=22 individual colonies from each group. Error bars represent SEM. Statistics were performed using two-tailed Student’s *t*-test.

To determine whether PCFUs maintain multi-lineage potential after multiple passages *in vitro*, differences in acinar, ductal, and endocrine lineage marker expression of individual colonies between the 1^st^ and 4^th^ generations were compared by microfluidic qRT-PCR. There was no observable difference in the morphology of cystic colonies (Fig. 3f) or gene expression (Fig. 3g) between the 1^st^ and 4^th^ generation colonies. These results demonstrate that PCFUs in the sorted FSC^mid-high^ population retain tri-potency over multiple generations *in vitro*.

### FSC^mid-high^ clusters contain tightly-bound individual cells resistant to enzyme dissociation *in vitro*

The small clusters in the FSC^mid-high^ fraction (referred to as FSC^mid-high^ clusters) are composed of multiple cells, so we sought to dissociate them into single cells for subsequent studies. Because we knew that FSC^mid-high^ clusters were resistant to collagenase (which was used to dissociate the pancreas prior to sorting), we employed trypsin. Remarkably, FSC^mid-high^ clusters remained intact even after a 1 hr incubation with 0.25% trypsin-EDTA (Fig. 4a), and retained their colony-forming ability compared to untreated FSC^mid-high^ clusters (Fig. 4b). Other enzymes, including Liberase and TrypLE, were tested with similar results (data not shown). FSC^mid-high^ clusters potentially resist dissociation due to strong cell-adhesion properties ^32^. To examine this, we performed immunofluorescence staining for classical tight junction markers and found that E-cadherin, TJP1 (ZO-1), and F11r (JAM-A) ^33,34^ were expressed at cell-cell interfaces of the FSC^mid-high^ clusters (Fig. 4c and Supplementary Fig. 3a). Interestingly, this was a feature found only in FSC^mid-high^ clusters; cystic colonies, formed after plating, expressed low levels of ZO-1 and JAM-A (Supplementary Fig.3b-c). Transmission electron microscopy (TEM) also revealed tight junctions at cell-cell boundaries of individual cells within FSC^mid-high^ clusters (Fig. 4d, yellow arrows), and each cell within the small cluster was single nucleated (Fig. 4d, nuclei labels). We counted the number of individual cells with nucleus per cluster and found that a cluster had an average of 8 cells (Fig. 4e and Supplementary Fig. 4). To visualize individual cells within the FSC^mid-high^ clusters in 3-dimensional (3D) space, we used serial block-face 3D scanning electron microscopy (3D-SEM). This confirmed the finding that FSC^mid-high^ clusters are comprised of multiple individual cells (Supplementary Movie). Taken together, these results demonstrate that FSC^mid-high^ clusters consist of individual single-nucleated cells that are tightly bound.

**Figure 4.**
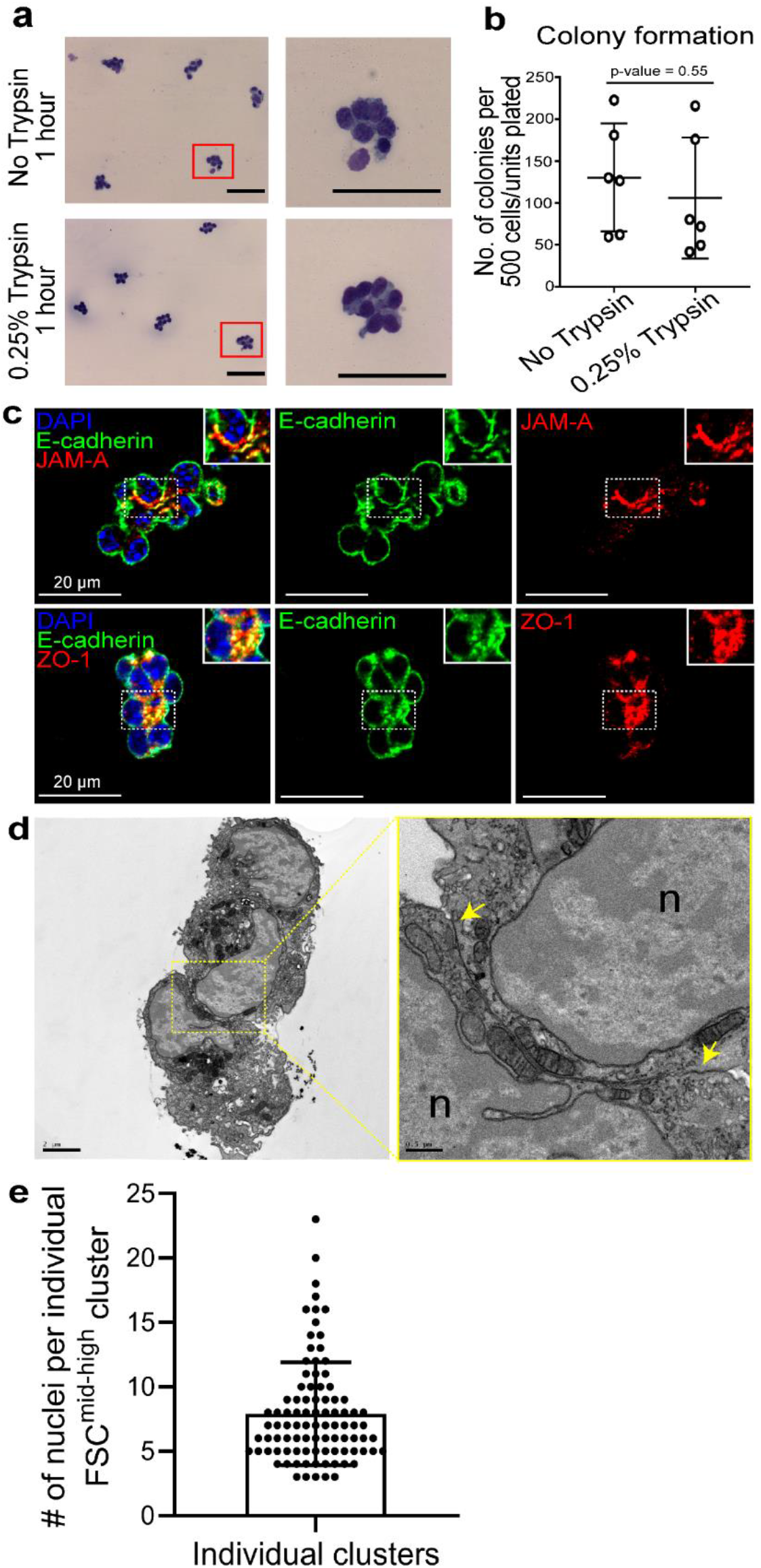
FSC^mid-high^ clusters exhibit strong cell-adhesion properties. (a) Sorted FSC^mid-high^ clusters was incubated with DMEM/F12 containing either DPBS (top) or 0.25% trypsin + EDTA (bottom) for 1 hr followed by Wright-Giemsa staining. Representative photomicrographs are shown. Scale bars=100 μm. (b) FSC^mid-high^ clusters treated with either 0.25% trypsin + EDTA or control vehicle for 1 hr and plated into the Matrigel/RSPO1 colony assay for 3 weeks. No significant difference between the resulting colonies from the control and trypsin-treated FSC^mid-high^ clusters. n=3 independent experiments with 2 biological replicates each. Statistics were performed using two-tailed Student’s t-test. Error bars represent SEM. (c) IF analysis of sorted FSC^mid-high^ clusters stained for tight junction markers which show expression at cell-cell interfaces for JAMA (top) or ZO-1 (bottom). All cells in a FSC^mid-high^ cluster stained positive for E-cadherin, indicating epithelial cell identity. Scale bars=20 μm. (d) Transmission electron microscopy (TEM) analysis of a FSC^mid-high^ cluster cross-section show two individual cells labeled with nuclei (n) bound together at cell-membranes with tight junctions (yellow arrows). Scale bar=2 or 0.5 μm, respectively. (e) Sorted FSC^mid-high^ clusters were stained by Wright-Giemsa and the number of nuclei in each cluster was manually quantified.

### Cell-tracing analysis reveals that FSC^mid-high^ clusters consist of cells with clonal potential that form individual cystic colonies

Despite the inability to dissociate FSC^mid-high^ clusters, we investigated whether colonies were derived from single or multiple cells. To test this, we employed a random X inactivation system and generated *Hprt^DsRed/+^* mice in which the gene for *Disocosoma* sp. red fluorescent protein (DsRed) replaced hypoxanthine guanine phosphoribosyl transferase (Hprt) on one X chromosome. In mammals, inactivation of one X chromosome takes place randomly in the pre-implantation female embryo ^35^ and is somatically inherited with extreme fidelity ^36^. As a result, hemizygous female *HprtD^sRed/+^* mice are cellular mosaics, and approximately half of their cells should express DsRed. As expected, fluorescence analysis and flow cytometry of pancreatic cells and splenocytes showed mosaic expression of DsRed in hemizygous female *Hprt^DsRed/+^* mice (Supplementary Fig. 5). For our purposes, it is important to emphasize that X inactivation is somatically heritable; the progeny of a DsRed^+^ cell will always be red and the progeny of a DsRed^-^ cell will always be devoid of the fluorescent signal.

The FSC^mid-high^ clusters from hemizygous female *Hprt^DsRed/+^* mice were cytospun onto slides, fixed, stained with DAPI, and analyzed (Fig. 5a). As expected, about half of the small clusters from FSC^mid-high^ fraction were mosaic (containing both DsRed^+^ and DsRed^-^ cells, 46.3%); in contrast, a quarter of small clusters were either fully labeled (DsRed^+^, 24.4%) or unlabeled (DsRed^-^, 29.3%; Fig. 5b-d). Given that mosaic cell clusters contained DsRed^+^ and DsRed^-^ cells, we predicted that plating mosaic cell clusters could form one of the following two types of colonies. First, mosaic colonies if multiple cells within a cluster proliferate to form the colony, showing non-clonality, or second, a fully labeled DsRed^+^ or unlabeled DsRed^-^ colony if an individual DsRed^+^ or DsRed^-^ cell within a cell cluster can self-renew and proliferate clonally.

**Figure 5.**
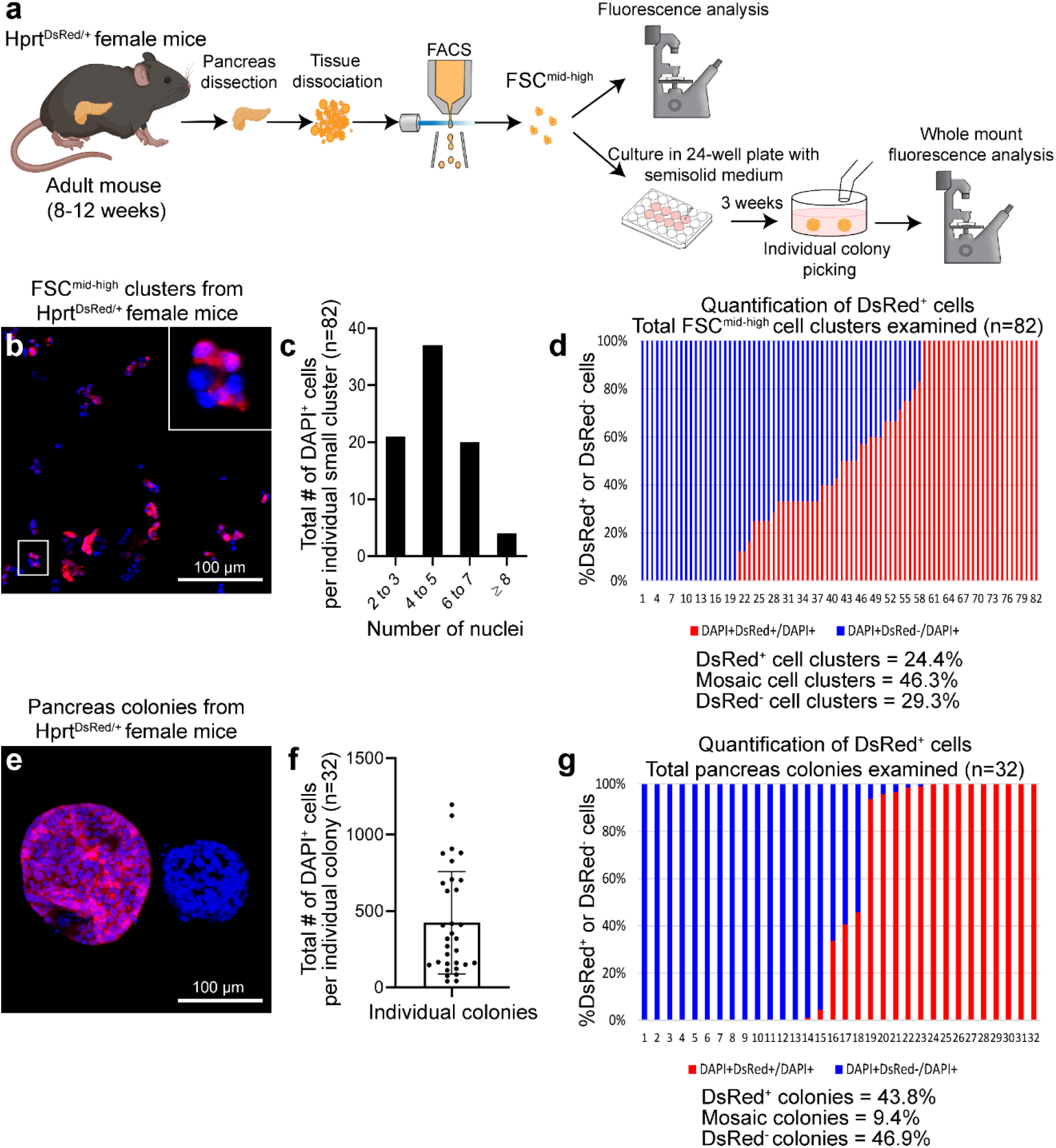
Cell-tracing analysis suggests colonies grown from FSC^mid-high^ clusters are clonal. (a) Schematic of experimental design. (b) Representative images of freshly-sorted FSC^mid-high^ clusters from Hprt^DsRed/+^ female mice. Nuclei were stained with DAPI (blue). The zoomed-in image of a cluster reveals a mixture of DsRed^+^ (red) and DsRed^-^ cells. Scale bar=100 μm. (c) Quantification of the number of nuclei per sorted FSC^mid-high^ cluster. n=82 clusters. (d) DAPI^+^ nuclei from each cluster were quantitated and the proportion between DsRed^+^ and DsRed^-^ cells was determined in each cluster. Each bar represents a FSC^mid-high^ cluster, n=82. FSC^mid-high^ clusters were categorized as DsRed^+^ (defined as >90% red cells; 24.4% among all clusters examined), mosaic (46.3%), or DsRed^-^ (defined as <10% red; 29.3%). (e) The sorted FSC^mid-high^ fraction from Hprt^DsRed/+^ female mice was plated into Matrigel/RSPO1 colony assay. Three weeks later, colonies were collected and imaged on a confocal microscope from top to bottom of each colony. Two representative cystic colonies, in the mode of Maximum Intensity Projection of all slices, are shown. Scale bar=100 μm. (f) The number of DAPI^+^ cells from each colony was quantitated. (g) Each bar represents a colony, n=32. Colonies were categorized as DsRed^+^ (defined as >90% red; 43.8% among all colonies examined), mosaic (9.4%), or DsRed^-^ (defined as <10% red; 46.9%).

Small clusters were plated into our Matrigel/RSPO1 colony assay and cultured for 3 weeks (Fig. 5a). Individual colonies were hand-picked, fixed, stained with DAPI, and examined under a confocal microscope using optical slice z-stacks (Fig. 5e). Total nuclei (DAPI^+^) and total DsRed^+^ cells were quantitated in 3D-reconstructed colonies; image analysis software was used to unbiasedly label DAPI^+^ cells as DsRed^+^ or DsRed^-^. We found that about half of the colonies were either fully labeled (43.8%, DsRed^+^) or unlabeled (46.9%, DsRed^-^), while only a few colonies were mosaic (9.4%, Fig. 5e-g). These data indicate that the majority of the colonies are clonally derived from a single cell within the originating small cluster. Interestingly, the sizes of individual colonies were not uniform (Fig. 5f), suggesting heterogeneous proliferative potential of the originating progenitor cells. Taken together, these cell-tracing experiments demonstrate that a single cell within a FSC^mid-high^ cluster possesses progenitor function, and we speculate that the neighboring cells may play other functions, such as regulating the proliferative potential.

### Droplet-based gene expression profiling reveals FSC^mid-high^ clusters express genes involved in cell-cell interactions, development, and cancer

To investigate gene expression patterns of the FSC^mid-high^ clusters, we performed droplet-based RNA sequencing with barcoding ^37^ (referred to as droplet RNA-seq) on both sorted FSC^mid-high^ and FSC^low^ control units. After running quality control (Supplementary Fig. 6a-c), a total of 1,125 FSC^mid-high^ and 598 sorted FSC^low^ units were analyzed (Fig. 6a). These two sets of data were merged, resulting in a total of 8 clusters (Fig. 6b-c and Supplementary Fig. 6d-f). Consistent with our prior finding that the parent CD133^high^CD71^low^ population is comprised of ductal cells ^23^, most of the clusters expressed ductal markers (*Sox9, Krt23, Krt17, Spp1*), except cluster 7 which expressed immune cell genes (Fig. 6b and Supplementary Fig. 6g-i). To validate the expression of lineage markers, we performed qRT-PCR on sorted FSC^low^ and FSC^mid-high^ fractions. As expected, both FSC^low^ and FSC^mid-high^ fractions expressed higher levels of ductal (*Sox9*) and lower levels of acinar (*Amylase2a*) and endocrine (*Insulin2*) markers, compared to unsorted total pancreatic cells (Supplementary Fig. 7a-c).

**Figure 6.**
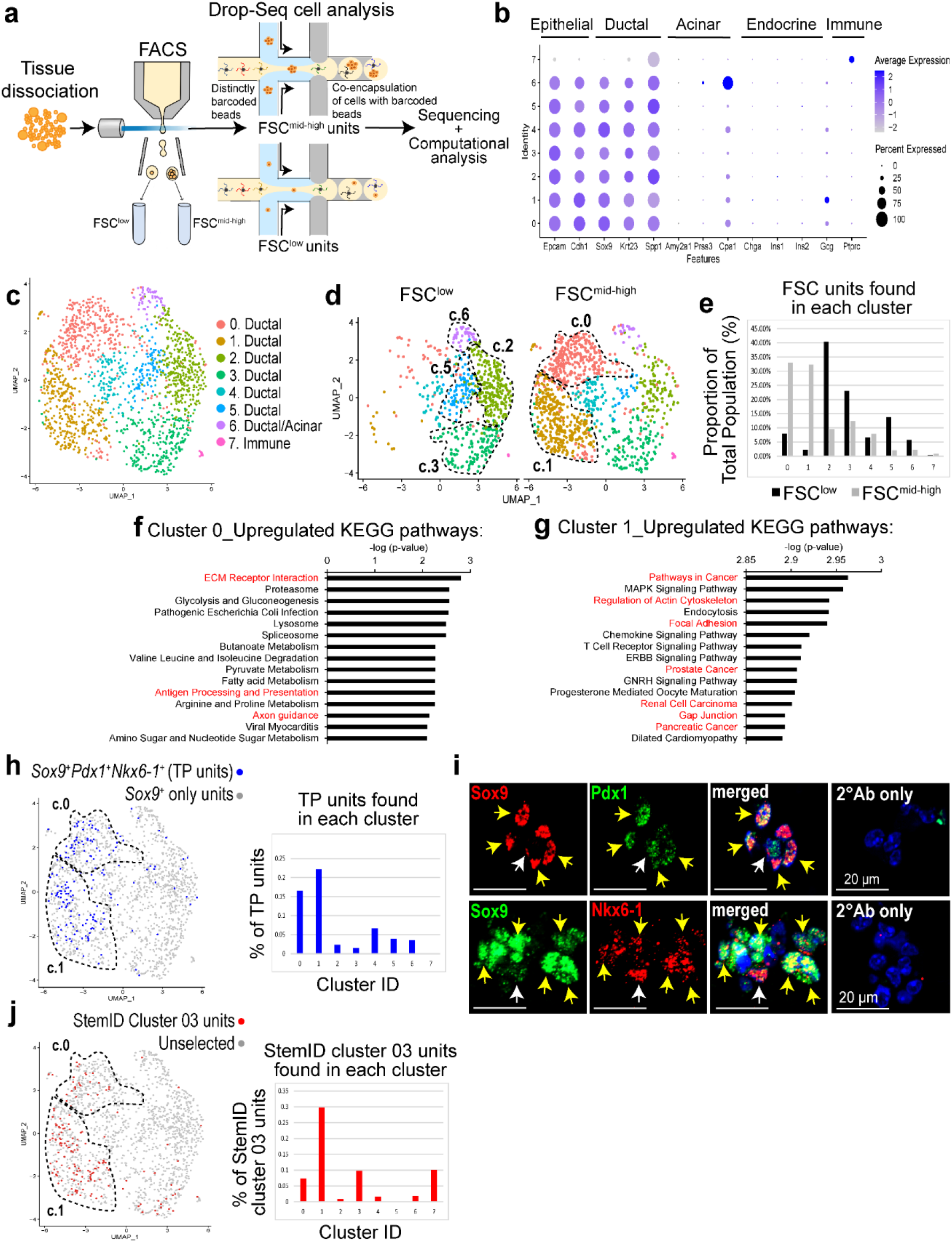
Transcriptomic analysis of FSC^mid-high^ clusters reveals a group of ductal cells with progenitor gene signature. (a) Schematic of experimental design for droplet RNA-seq (Drop-Seq) analysis of FSC^low^ and FSC^mid-high^ fractions. After quality control was run, datasets from FSC^low^ (n=598) and FSC^mid-high^ (n=1,125) populations were combined (total 1,723) and clustered using Seurat. (b) Dot plot of the average scaled expression (measured by average Pearson residual) of canonical markers for epithelial, pancreatic, and immune cell types, plotted against cluster identity. (c) Uniform manifold approximation and projection (UMAP) visualization of 8 identified clusters (0 to 7) of the combined datasets. (d) UMAP visualization, separating FSC^low^ and FSC^mid-high^ populations. (e) Proportion of FSC^low^ units and FSC^mid-high^ units found in each cluster from total population. (f-g) Gene set enrichment analysis (GSEA) of differentially expressed (DE) genes found in Cluster 0 and Cluster 1 to identify biological relevant pathways using GO (Supplementary Dataset 2-3) and KEGG. Adjusted p-value < 0.05. (h) Units that co-express the three genes *Sox9, Pdx1*, and *Nkx6-1* are identified as triple-positive (TP) units and labeled in the UMAP as blue dots (left). Percentage of TP units per cluster (right). (i) IF analysis of FSC^mid-high^ clusters shows Sox9^+^ ductal cells co-expressing Pdx1 or Nkx6-1 markers (yellow arrows). The white arrows indicate Sox9^-^Pdx1^+^ cells (upper panel) or Sox9^-^Nkx6-1^+^ cells (lower panel). Scale bar=20 μm. (j) Aggregated dataset was first processed using StemID algorithm and the resulting StemID cluster 03 was identified with the highest StemID score (Supplementary Fig. 10). The StemID cluster 03 (red dots) was then back-fitted in the Seurat UMAP clusters 0 to 7 (left). Percentage of StemID cluster 03 units was highest in UMAP cluster 1 (right). Abbreviations: GO, Gene Ontology; KEGG, Kyoto Encyclopedia of Genes and Genomes.

Further analysis of the seven ductal clusters revealed that clusters 0 and 1 were most enriched in the FSC^mid-high^ fraction, while clusters 2, 3, 5, and 6 were most enriched in the FSC^low^ fraction (Fig. 6d-e). Cluster-specific differentially-expressed genes (Supplementary Dataset 1) were further analyzed by gene set enrichment analysis (GSEA) using Gene Ontology and KEGG molecular signature databases (Supplementary Dataset 2 and 3). Common among clusters 0 and 1, which were enriched in the FSC^mid-high^ fraction, were up-regulated pathways involved in cell-cell interactions/signaling and organ development (Fig. 6f-g and Supplementary Fig. 8a-b). Interestingly, various cancer pathways were significantly up-regulated in cluster 1, but not cluster 0 (Fig. 6f-g). In sharp contrast, most pathways that were up-regulated among clusters 2, 3, 5, and 6 were involved in metabolism, although pathways in cellular differentiation and proliferation were occasionally noted in cluster 2 (Supplementary Fig. 8a-b). These results suggest functional diversity and heterogeneous nature of ductal cells.

As previously mentioned, embryonic pancreatic MPCs are responsible for expansion and differentiation into all three main pancreatic lineages and they express the transcription factors *Sox9, Pdx1*, and *Nkx6-1* ^30,31^. We hypothesized that clusters 0 and 1 could be enriched with units that co-express these genes, which we termed triple-positive cell units (TP units). Differentially-expressed genes in TP units are presented in Supplementary Dataset 1. Indeed, clusters 0 (16.5%) and 1 (22.1%) contained the higher proportions of TP units compared to clusters 2 - 7 (7% or less) (Fig. 6h). Immunofluorescence staining of small clusters from FSC^mid-high^ fraction confirmed individual nuclei that co-expressed either Sox9 and Pdx1 or Sox9 and Nkx6-1 proteins (Fig. 6i). Further analysis of other established MPC markers ^38^ revealed that clusters 0 and 1 express *Hnf1b, Hes1, Foxa2, Tead1, Nkx2-2, Rbpj, Glis3, Nr5a2, Gata6, Yap1, Taz*, and *Myc*, but not *Ptf1a* and *Gata4* (Supplementary Fig. 9). Taken together, these results demonstrate that clusters 0 and 1 are enriched for TP units that resemble MPCs.

To further explore the idea of clusters 0 and 1 being enriched for progenitor cells, we employed the StemID algorithm ^39^, a mathematical tool that has been successfully used to identify rare stem and progenitor cells (i.e. hematopoietic and intestinal stem cells) within a heterogenous population of cells. StemID independently reclustered our dataset into 18 new clusters (Supplementary Fig. 10a), and branch point analysis rendered a lineage tree showing differentiation trajectories (Supplementary Fig. 10b). The StemID score for each cluster was calculated ^39^, and StemID Cluster 3 had the highest StemID score, which indicated the highest level of multipotency among all units detected in this dataset (Supplementary Fig. 10c-h). The StemID cluster 3 was composed of 167 units (differentially-expressed genes are presented in Supplementary Dataset 1), which were enriched in the FSC^mid-high^ fraction compared to FSC^low^ fraction (Supplementary Fig. 10i). Importantly, the 167 units found in StemID cluster 3 were enriched in cluster 1 (29.7%) compared to all other clusters (10% or less) in our original clustering (Fig. 6j), which suggests that cluster 1 is most enriched for progenitor-like cells. Notably, several differentially-expressed genes were found overlapping between cluster 1, TP units, and StemID cluster 3 (Supplementary Fig. 11), suggesting commonality among these cells. Taken together, our transcriptomic analyses highlighted cluster 1 (primarily FSC^mid-high^ units) as being most enriched for progenitor-like cells.

### FSC^mid-high^ clusters are enriched after ablation of acinar cells *in vivo*

As demonstrated in Fig. 4, the ductal FSC^mid-high^ clusters remained intact after prolonged treatment with enzymes *in vitro*. This raised the possibility that the FSC^mid-high^ clusters may survive in adult mice after severe damage to acinar cells, which release digestive enzymes *in situ* ^40^. To test this, we used ElaCreERT2 mice with floxed-Diphtheria toxin receptor (DTR) to induce acinar cell death ^41^. Adult ElaCreERT2;R26^DTR/DTR^ mice were injected with tamoxifen (TAM) to induce DTR expression in acinar cells, then were treated with diphtheria toxin (DT) (Fig. 7a). Three days after the last dose of DT, we found that body weight was similar between control and injured groups (Supplementary Fig. 12a). However, pancreata from injured groups were significantly smaller in size due to significant loss of acinar cells compared to controls (Fig. 7b-c and Supplementary Fig. 12b-c). Also, observable areas of translucent patches, indicative of pancreatic edema, in both the head and tail of the pancreas were found in injury groups compared to controls (Fig. 7b-c). These results demonstrate an effective mouse model to induce severe acinar cell injury *in vivo*, which allowed us to subsequently study its effects on leftover ductal cells, including the FSC^mid-high^ clusters.

**Figure 7.**
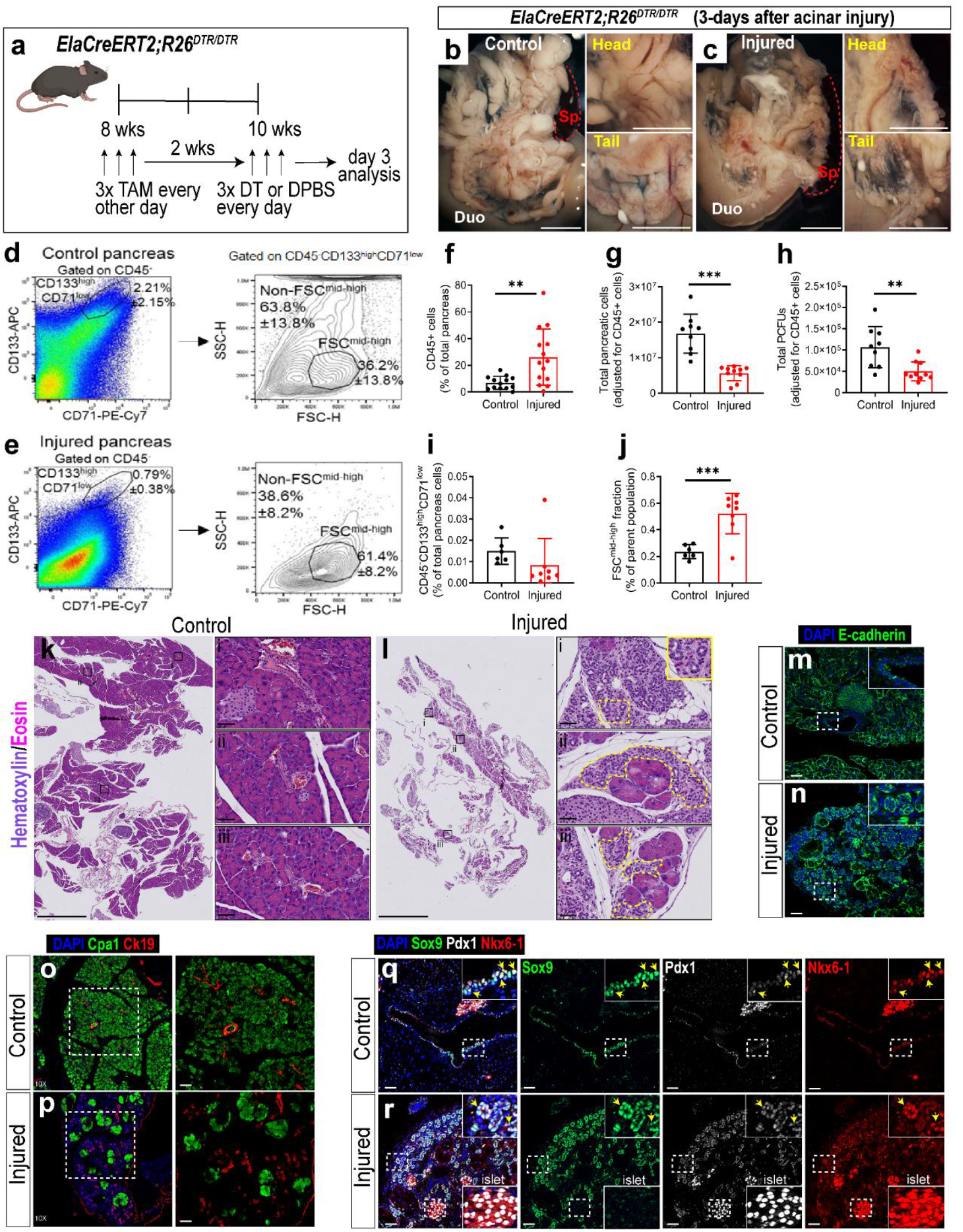
FSC^mid-high^ clusters can survive *in vivo* acinar injury conditions, which induce the formation of ductal rosettes that contain TP cells. (a) Scheme for pancreatic acinar injury model. ElaCreERT2;R26^DTR/DTR^ mice were injected with tamoxifen (TAM) to induce the expression of diphtheria toxin receptor (DTR) in acinar cells, followed by injections of diphtheria toxin (DT) to destroy acinar cells. (b-c) Representative brightfield images of control and injured pancreas 3 days after the last DT injection. Abbreviations: Duo, duodenum; Sp, spleen; Head, dorsal pancreas; Tail, ventral pancreas. Red dotted lines outline the spleen. Scale bars=1 mm. (d-e) Flow cytometry analysis of the parent CD133^high^CD71^low^ ductal population after gating on CD45-negative cells, followed by light scattered analysis to identify the percentage of FSC^mid-high^ subpopulation compared to non-FSC^mid-high^ subpopulation. (f) The pan-leukocyte marker CD45 showed increased infiltration of CD45^+^ cells in the injured pancreata compared to controls. **p<0.01, n=13-14. (g) The total number of cells per pancreas, after adjusting for % CD45^+^ cells, was reduced in the injured pancreata compared to controls. ***p<0.001, n=9-10. (h) Unsorted pancreas cells from control and injured pancreata were plated in Matrigel/RSPO1 colony assay to determine colony-forming efficiency, and subsequently the total number of PCFUs per pancreas was calculated and presented. **p<0.01, n=9-10. (i) Percentage of the parent CD133^high^CD71^low^ ductal population after gating on CD45-negative cells. n=6-8. (j) Percentage of FSC^mid-high^ subpopulation compared to non-FSC^mid-high^ subpopulation. ***p<0.001, n=6-8. Statistics were performed using two-tailed Student’s *t*-test. (k-l) Representative Hematoxylin and Eosin (H&E) staining images of samples 3 days after injury. Insets for control pancreas (k.i - k.iii) show normal intercalated ducts, whereas, insets for injured pancreas (l.i - l.iii) show rosette structures near acinar cells. Dashed lines represent zoomed-in or outlined rosettes. Scale bars=2.5 mm. (m-n) IF analysis of pancreas samples for epithelial marker E-cadherin. Scale bars=50 μm. (o-p) IF analysis of pancreas samples co-stained with acinar (Cpa1) and ductal (CK19) markers. Images on left are 10X magnified. Scale bars=50 μm. (q-r) IF analysis of pancreas samples 3 days after injury co-stained with Sox9, Pdx1, and Nkx6-1. White-dashed boxes zoomed into insets of ductal cells co-expressing the three markers (yellow arrows). Insets of islets confirm co-expression of Pdx1 and Nkx6-1 with no Sox9. Scale bars=50 μm.

Immune cells are known to migrate to the pancreas after severe damage ^42^. We therefore measured the infiltration by CD45^+^ leukocytes using flow cytometry (Fig. 7d-e and Supplementary Fig. 13). There was a significant increase in the percentage of CD45^+^ cells in the injured pancreas compared to control (Fig. 7f). Because of this significant leukocyte infiltration, we subtracted these cells when calculating the total number of pancreatic cells in each mouse, which was lower in the injured pancreas compared to controls (Fig. 7g). Next, to determine if acinar cell injury affects the remaining pancreas cells and their colony-forming capability, we plated unsorted cells from injured pancreata and controls into our Matrigel/RSPO1 colony assay. Using the total number of pancreatic cells and colony-forming efficiencies in each mouse to calculate, we found that the total number of PCFUs per pancreas was significantly reduced in the injured samples compared to controls (Fig. 7h). This result suggests that at least some PCFUs are lost after pancreas injury. To probe this further, we found that the proportion of parent CD133^high^CD71^low^ cells among the total pancreatic cells was not significantly different (Fig. 7i), suggesting that the parent ductal cells were maintained. However, the proportion of the FSC^mid-high^ sub-population among the other FSC sub-populations was significantly increased in injured pancreata compared to controls (Fig. 7j). These results were similar between male and female cohorts (Supplementary Fig. 14). Together, these data demonstrate that the FSC^mid-high^ clusters, compared to non-FSC^mid-high^ clusters, preferentially survive under *in vivo* acinar cell injury.

### Ductal rosette structures in injured pancreas samples contain TP cells and undergo proliferation 14-days after injury

Given that our acinar cell injury model results in significant acinar cell loss, we hypothesized that ducts, which contain FSC^mid-high^ clusters, could become proliferative and contribute to tissue regeneration. We first confirmed acinar cell loss in injured samples using H&E staining (Fig. 7k-l). During this analysis, we observed many cell rosette structures near remaining acinar cells in injured samples (Fig. 7l, i-iii insets), which we confirmed to be epithelial (E-cadherin^+^) (Fig. 7m-n). Interestingly, these rosette structures were also found in an acute acinar injury model (Supplementary Fig. 12d, i-iii insets), in which we used a lower DT dose ^41^. To test whether the epithelial rosettes express markers for acinar-to-ductal cell metaplasia (ADM) ^43^, which usually occurs after acinar cell injury, we examined the co-expression of CK19 and Cpa1 (Fig. 7o-p) as well as CK19 and Amylase (Supplementary Fig. 12e-f). Epithelial rosettes expressed CK19 only, suggesting that epithelial rosettes were not undergoing ADM 3-days after the last DT injection. Next, because the FSC^mid-high^ clusters can express Sox9, Pdx1, and Nkx6.1 (Fig. 6j), we examined whether Sox9^+^/Pdx1^+^/Nkx6.1^+^ triple-positive cells (referred to as TP cells) can be found in the control and injured ducts. In control mice, TP cells were found in main, small interlobular, and intercalated ducts (Fig. 7q; upper panel and Supplementary Fig. 12g). However, at 3-days post-injury the TP cells were found in ductal rosettes (Fig. 7r; lower panel). Next, we used EdU labeling following acinar injury to determine if ductal rosettes may contribute to repair by proliferating (Fig. 8a), but very few proliferating cells were found in ductal rosettes 3-days post-injury (Fig. 8b-c). One potential reason for the lack of proliferating cells in ductal rosettes could be that early injury conditions inhibit the activation of proliferation, which is known to occur in neurons after brain injury ^44^. Thus, we analyzed injured pancreata 14-days post-injury (Fig. 8d), which showed a continued reduction in pancreas weight (Supplementary Fig. 15a-c) and severe loss of acinar cells. Notably, rosette structures were still nearby the acinar cells (Supplementary Fig. 15d-e). Immunofluorescence analysis confirmed the continued presence of TP cells in ductal rosettes (Fig. 8e-f and Supplementary Fig. 15f-k). At this time point, an increased percentage of Sox9-positive ducts co-labeled with EdU and/or Ki67 (another marker for proliferation) among injured sample rosettes compared to control pancreata was found (Fig. 8g-k and Supplementary Dataset 4). Taken together, these findings show that severe ablation of acinar cells in the adult mouse pancreas leads to the formation of ductal rosette structures that contain TP cells, which undergo proliferation 14-days post-injury, potentially contributing to pancreas organ regeneration *in vivo*.

**Figure 8.**
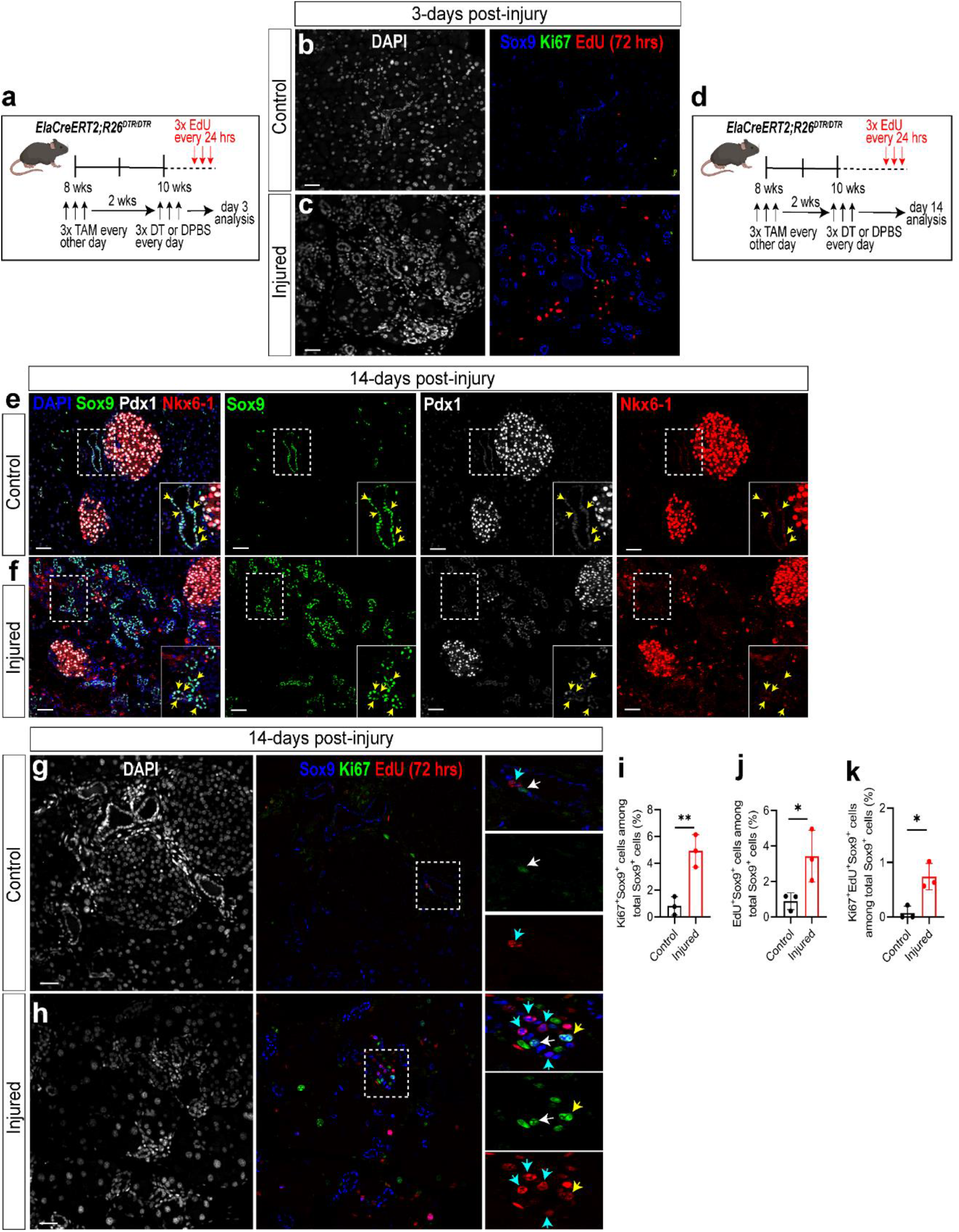
Ductal rosettes contribute to pancreas proliferation 14-days after acinar injury. (a) Experimental scheme for a 72-hr EdU labeling of pancreas 3 days after the last dose of DT injection to induce acinar injury. (b-c) IF analysis of Sox9 (ductal marker), Ki67 (proliferation marker), and EdU label (S-phase cell cycle analog) in pancreas samples 3 days after acinar injury. Sox9^+^ ductal cells were not proliferating. Scale bars=50 μm. (d) Experimental scheme for a 72-hr EdU labeling 14 days after the last dose of DT injection. (e-f) IF analysis of pancreas samples 14 days after injury co-stained with Sox9, Pdx1, and Nkx6-1. White-dashed boxes were zoomed into insets of ductal cells which show co-expression of the three markers (yellow arrows; termed TP cells). Scale bars=50 μm. (g-h) IF analysis of Sox9^+^ ductal cells co-expressing Ki67 and/or EdU label in pancreas samples 14 days after acinar injury. Sox9^+^ cells co-expressing Ki67 (white arrows) or EdU (light blue arrows) or both (yellow arrows) are shown. Scale bars=50 μm. (i-k) Quantification of the percentage of Sox9^+^ ductal cells co-expressing Ki67 (i), EdU (j) or both (k) from control and injured pancreas samples. *p<0.05, **p<0.01, n=3.

## Discussion

In this study, we fractionate CD133^high^CD71^low^ ductal cells (Fig. 1) from adult murine pancreas based on forward and side scatters in flow cytometry and discovered a previously unknown ductal cell cluster (termed FSC^mid-high^) that exhibits progenitor cell properties, which include the ability to self-renew robustly (~100,000 fold over 9 weeks; Fig. 3) and differentiate into the three pancreas cell lineages *in vitro* (Fig. 2). We discovered that FSC^mid-high^ clusters are composed of individual cells that express tight junction markers at the cell-cell interfaces, which may explain their resistance to enzymatic dissociation *in vitro* and *in vivo* (Fig. 4 & 7). Furthermore, individual cells within the FSC^mid-high^ clusters expand clonally (Fig. 5), which may be similar to the previously-reported structural proliferative units identified in the pancreas of post-natal mice using an X-linked reporter gene ^45^. Adult FSC^mid-high^ clusters express embryonic MPC markers such as Sox9, Pdx1, and Nkx6-1, which we refer to as TP cells (Fig. 6). Importantly, we located TP cells within the ducts from normal and injured adult pancreata (Fig. 7). Upon severe acinar cell injury, ductal rosettes that are comprised of TP cells become proliferative, potentially contributing to pancreas organ recovery (Fig. 8). In all, our data support the existence of a sub-set of ductal cells (i.e., FSC^mid-high^ clusters) with progenitor cell properties that may play roles in pancreas organ homeostasis and/or disease initiation/progression.

A finding of the current study is the demonstration of heterogeneity of the adult ductal cells. Historically, cells in the duct, acinar, and endocrine cell compartments have been largely considered homogeneous. However, increasing evidence in recent years suggest heterogeneity in all three pancreatic compartments ^39,46,47^. We have previously shown that CD133^+^ total ductal cells comprise 13.1 ± 4.3% of the pancreatic cells in adult mice ^25^. The CD133^high^CD71^low^ ductal cells comprise 2.4 ± 1.9% of the total pancreatic cells ^23^. In this study, we find that the CD133^high^CD71^low^FSC^mid-high^ fraction is only 0.12 ± 0.04% of the total pancreatic cells. With such a small proportion of cells, researchers who use pan-ductal markers such as Sox9 ^14^ or HNF1b ^13^ to conduct *in vivo* lineage-tracing experiments are limited to observe a very minor contribution of progenitor cells. Specifically, in the homeostatic state a minor ductal population may not be identified above the ‘noise’, missing significant contributions to pancreatic regeneration and potentially tumorigenesis. This is supported by a recent lineage-tracing study showing that a small population of Ngn3-expressing ductal cells give rise to somatostatin and insulin expressing cells in the adult mice ^18^.

Our discovery that FSC^mid-high^ clusters retain colony-forming ability even after 1 hour of trypsin digestion *in vitro* (Fig. 4) was unexpected. We found only one other example in the literature for the pituitary gland from which small clusters of cells are also resistant to trypsin digestion *in vitro* while retaining the ability to form colonies that express pituitary gland hormones ^48^. The biological significance of the clustering of normal cells has not been elucidated; we speculate that these cells may contribute to the *in vivo* niche, similar to the relationship of a Paneth to intestinal stem cell which increases stem cell colony formation *in vitro* ^49^. Our Gene Ontology (GO) analysis show that cluster 0 is enriched in pathways involved in regulation of ECM-receptor interaction, neuron differentiation, vascular development, blood vessel morphogenesis, and regulation of axonogenesis (Supplementary Fig. 8a), which may suggest a role in recruiting neurons and blood vessels to support tissue homeostasis ^50^. On the other hand, cluster 1 was primarily enriched in pathways involved in response to growth factor, cell division, regulation of growth, and microtubule cytoskeleton organization (Supplementary Fig. 8a). This may suggest that cluster 0 play a niche role in supporting progenitor cells ^51^ contained in cluster 1; however, this possibility requires further investigation.

Another aspect for the tight cluster could be similar to a function observed in tumor cells to enhance movement and metastasis ^52^. Consistent to this idea, some of the pathways enriched in the FSC^mid-high^ clusters involve cell migration (Supplementary Data Sets 2 and 3). We speculate that after injury, FSC^mid-high^ clusters that contain TP units survive enzyme digestion and migrate to where they are needed for organ regeneration.

The pancreas has a low cell turnover rate and most cells are non-proliferative ^53–55^. During insults such as pancreatitis, various pancreatic cells become activated to initiate repair and regeneration ^56^. Consistent with this body of prior work, we show that the ductal rosettes express proliferation markers on day 14 after injury (Fig. 8). After acute pancreatic injury, acinar cells often undergo ductal metaplasia as part of the regeneration process ^57^. Given that we did not observe ductal rosettes simultaneously expressing ADM markers day 3 or 14 post-injury (Fig. 7 and 8), it is unlikely that ductal rosettes after injury is mediated through the ADM process. Instead, these ductal clusters may harbor their own regenerative potential that works independently or in concert with other regenerative mechanisms.

We find that pancreatic cancer-related gene sets were enriched in gene expression patterns identified from the FSC^mid-high^ fraction, suggesting that progenitor features are associated with tumorigenesis ^58^. Other groups have suggested distinct acinar versus ductal tumor-initiating cells for basal-like or classical subtypes of pancreatic ductal adenocarcinoma ^11,59^. Our results raise the possibility that the FSC^mid-high^ clusters may represent a population of cells that posess survival advantage and are amenable to transformation in the ductal compartment. Thus, the adult ductal clusters, which express embryonic MPC markers as described in this report, have implications in both regenerative medicine and tumorigenesis.

In summary, we have identified multi-cellular units in the adult mouse pancreas capable of self-renewal and differentiation *in vitro* that are resistant to severe acinar cell ablation *in vivo*. These cell clusters express conventional ductal markers but are enriched for genes involved in pathways known for embryonic pancreatic multipotent progenitor cells and pancreatic cancer. Our results implicate these ductal progenitor-like cells in both regenerative medicine and tumorigenesis.

## Materials and Methods

### Mice

C57BL/6J (B6) mice (The Jackson Laboratory, Bar Harbor, ME) (8-12 week-old, both sexes) were used in most experiments unless specified otherwise. Transgenic mice harboring ElaCreERT2 and R26^DTR^ alleles were as reported previously ^41^. Congenic Ela-CreERT2;R26^DTR/DTR^ mice were produced by backcrossing to B6 mice to yield >99.99% B6 genetic background; 8-12 week-old male and female mice were used for injury studies. FVB.129S1-Hprt^tm1(CAG-DsRed)Mnn/COH^ (*Hprt*^DsRed/+^) mice were generated in-house using gene targeting vectors and embryonic stem (ES) cell clone selection strategies. The coding sequence (cds) of the T1 variant of *Discosoma* sp. red fluorescent protein (DsRed), also known as DsRedExpress ^60^, was obtained from Clontech and knocked-in at the X-linked hypoxanthine guanine phosphoribosyl transferase (*Hprt*) locus, as previously described ^61^, except: (1) the DsRed cds was used in place of the Cre cds, and (2) *loxP* sites flanked the neo selection cassette. Neomycin (neo)-resistant clones were genotyped using Southern blots to identify putative *Hprt*^DsRed/+^ 129S1 recombinants. *Hprt*D^sRed/+^ male chimeras were mated to 129S1/SvImJ (129S1) females to confirm germline transmission of the transgene. The neo selection cassette was removed by breeding with 129S1 Cre deleter mice ^61^ to produce the final knock-in line. Subsequently, Hprt^DsRed/+^ mice were bred and phenotyped using visual observation of red fluorescence with a TRITC/Cy3 filter set for excitation 454 ±15 nm and emission 620 ± 30 nm wavelength. *Hprt*D^sRed/+^ 129S1 mice were back-crossed to FVB/NJ mice for >10 generations to produce an *Hprt*^DsRed/+^ 129S1.FVB/NJ congenic strain to improve fluorescence imaging in live mice and increase average litter size. *Hprt*^DsRed/+^ female (8-12 week-old) chimeras were used for FSC^mid-high^ cluster and colony analysis. All mice were maintained under specific pathogen-free conditions. Animal experiments were conducted according to the Institutional Animal Care and Use Committee at the City of Hope.

### Dissociation of pancreas

Murine pancreata were dissected, cleared of fat tissue under a dissecting microscope, and rinsed three times in cold Dulbecco’s phosphate-buffered saline (DPBS) containing 0.1% bovine serum albumin (BSA), 100 Units (U)/mL penicillin, and 100 μg/mL streptomycin; this wash solution is referred to as DPBS/BSA. Pancreata were minced in a dry petri dish on ice using spring scissors for 3 min or until finely minced. Tissue was transferred to a 50 mL conical tube and resuspended in DPBS/BSA containing collagenase B (2-4 mg/mL) (Roche, Mannheim, Germany) and DNase I (2,000 U/mL) (Calbiochem, Darmstadt, Germany). Tissue was incubated at 37°C for 16 min; the tissue was swirled every 2-3 min and gently passed through a 16G syringe every 8 min. Cells were washed in cold DPBS/BSA containing 2,000 U/mL DNase I and successively passed through 100 μm and 40 μm mesh filters (BD Biosciences, San Jose, CA) to yield a single-cell suspension.

### Sorting and flow cytometry analysis

Sorting was performed similarly to our previous publication ^23^. Briefly, dissociated pancreatic cells/units were incubated with anti-mouse CD16/32 (10 μg/mL; BioLegend, San Diego, CA) for 5 min on ice to reduce non-specific binding. Biotin-conjugated anti-mouse CD133 (clone 13A4; 5 μg/mL; eBioscience, San Diego, CA) and phycoerythrin/Cy7 (PECy7)-conjugated anti-mouse CD71 (clone RI7217; 5 μg/mL; BioLegend, San Diego, CA) antibodies were added. Cells/units were incubated for 20 min on ice, washed twice, treated with streptavidin-labeled allophycocyanin (APC) (2 μg/mL BioLegend) for 15 min on ice, washed twice, and resuspended in DPBS/BSA/DNase I containing DAPI (0.2 μg/mL). Control antibodies were biotin-conjugated rat immunoglobulin (Ig) G1 (5 μg/mL; eBioscience, San Diego, CA) and PE/Cy7-conjugated rat IgG1 (5 μg/mL; BioLegend, San Diego, CA). Flow cytometry data were collected using Fortessa LSRII (Becton Dickinson, San Jose, CA) or Attune NX Cytometer (Thermo Fisher, Waltham, MA), with sorting performed on an Aria special order research product (SORP) (Becton Dickinson, San Jose, CA). All data were analyzed using FlowJo software (TreeStar, Ashland, OR). Analyses included initial gating of forward (FSC) and side (SSC) scatters to exclude debris. In sorting experiments, doublets were excluded by gating out high pulse-width cells; live cells were selected by DAPI-negative staining.

### Colony Assay

Sorted cells/units were resuspended at a density of 500 cells/units per well per 0.5 mL for the Matrigel/RSPO1 colony assay or 8.0×10^3^ cells/units per well per 0.5 mL for the laminin hydrogel assay as described previously ^25^. Culture media contained DMEM/F12 media, 1% methylcellulose (Shin-Etsu Chemical, Tokyo, Japan), 50% conditioned media from mouse embryonic stem cell-derived pancreatic-like cells ^62^, 5% fetal calf serum (FCS), 10 mmol/L nicotinamide (Sigma, St. Louis, MO), 10 ng/mL human recombinant activin B (R&D Systems, Minneapolis, MN), 0.1 nmol/L exendin-4 (Sigma), and 1 ng/mL vascular endothelial growth factor-A (VEGF) (Sigma). When indicated, either 5% (vol/vol) Matrigel plus 750 ng/mL mouse recombinant RSPO1 (R&D) (referred to as the Matrigel/RSPO1 colony assay in this study) or 100 μg/mL of laminin hydrogel ^25^ (referred to as the laminin colony assay in this study) was added to media to generate colonies. Cells/units were plated in 24-well ultra-low protein-binding plates (Corning, New York, USA) and incubated in a humidified 5% CO2 atmosphere at 37°C. Colonies grown in Matrigel/RSPO1 or laminin assay were counted 3 weeks or 10 days after plating, respectively. Colony-forming efficiency, or percent (%) PCFU, was determined by counting the total number of colonies per well after 2-3 weeks divided by the number of cells/units plated on day 0.

### Micro-manipulation of single cells/units or colonies

Sorted cells/units were placed in Matrigel/RSPO1 colony assay at a concentration of 1.0×10^3^ cells/units per mL ^63^. Single cells/units were visualized under a phase-contrast microscope, individually lifted using a fine Pasteur pipet with a ~30 μm diameter opening, and placed into the standard colony assay at a concentration of 1 cell/unit per well in a low-attachment Nunclon 96-well plate (ThermoFisher). Colonies were counted 2 weeks after plating. For micro-manipulation of single colonies, colonies grown in the 24-well plate described above were individually lifted using a 10-μL Eppendorf pipette set to 2 μL under direct microscopic visualization.

### Serial colony dissociation and replating

Warmed DPBS/BSA (1 mL/well) was added to each colony-containing well of a 24-well plate in standard Matrigel/RSPO1 colony assay. Colonies were collected in a 50 mL conical tube, washed, resuspended in 10 mL of 2-4 mg/mL collagenase B, incubated for 15 min at 37°C with mixing every 5 min, and washed in DPBS/BSA. Subsequently, colonies were treated with 20 mL of 0.25% (wt/vol) trypsin-EDTA, incubated for 3 min at 37°C, and pipetted thoroughly. Warmed FCS (4 mL) was added to stop trypsin digestion. Cells were washed in DPBS/BSA and kept at room temperature. A portion of the cells was mixed with 0.02% (wt/vol) trypan blue; the concentration of live cells (i.e. trypan blue-negative cells) was determined using a hemocytometer. For replating experiments, the final cell suspension was mixed with Matrigel and RSPO1-containing medium as indicated above. The total number of PCFUs was calculated by multiplying the previous dilution factor(s) with the number of colonies, per well, in the present culture.

### Cytospin and Wright-Giemsa Staining

The Shandon Cytospin 4 centrifuge system (ThermoFisher, Waltham, MA) was used to deposit a thin layer of cells onto Superfrost Plus microscope slides (ThermoFisher). A total of 2,500 cells/units from each sorted sub-population were aliquoted into 150 μL FCS, added to the EZ single Cytofunnel, and spun at 1,400 rpm for 5 min. For trypsin digestion, sorted FSC^mid-high^ cells/clusters were placed in low-binding 96-well plates at 2,500 units per well with 0.25% trypsin-EDTA and incubated at 37°C for 1 hr. The reaction was stopped by adding 100% FCS, and the cells/clusters were spun onto slides. Slides were removed, air-dried for 5 min, and incubated with ice-cold 100% methanol for 5 min to fix cells/clusters. Subsequently, cells/clusters were stained with 10% modified Wright-Giemsa solution (Sigma) for 30 min, washed with water, and imaged using a Zeiss Observer II (ZEISS, Oberkochen, Germany).

### Conventional or Microfluidic Quantitative Reverse Transcription-Polymerase Chain Reaction (qRT-PCR)

For conventional and microfluidic qRT-PCR analyses, the same procedures were employed as reported ^25^. Microfluidic qRT-PCR was performed using the BioMark™ and 48.48 Dynamic Array system (Fluidigm, South San Francisco, CA). Single colonies were individually lifted under direct microscopic visualization using a 10 μL Eppendorf pipette, collected in reaction buffer (10 μL), and pre-amplified (12 or 18 cycles for Matrigel/RSPO1- or laminin-grown colonies, respectively) according to the manufacturer’s instructions (Fluidigm). Amplified cDNA was loaded onto a 48.48 Dynamic Array using a NanoFlex integrated fluidic circuit (IFC) controller (Fluidigm). Threshold cycle (Ct), a measure of fluorescence intensity, was determined using BioMark PCR analysis software (Fluidigm) and expressed as delta Ct of the housekeeping gene beta-actin. All experiments were performed with negative (water) and positive (adult C57BL/6J pancreatic cells) controls. Taqman probes used in this study are listed in Supplementary Table 1.

### Droplet-based RNA-sequencing

FSC^mid-high^ and FSC^low^ sub-populations were sorted, counted, and diluted to the manufacturer’s recommended concentration in DPBS supplemented with 0.1% BSA. FSC^mid-high^ and FSC^low^ units were captured on a 10x Chromium device using a 10X V3 Single Cell 3’ Solution kit (10x Genomics, Chromium Single Cell 3’ Regent 00kit V3 Chemistry, Cat. PN-1000092). All protocols were performed following the manufacturer’s instructions. Final sequencing libraries were analyzed on a High Sensitivity DNA Chip (Agilent, Cat 5067-4626) to determine library size; final library concentrations were determined using a Qubit High Sensitivity DNA Assay Kit (ThermoFisher). Libraries were sequenced using the paired end setting of 101-101 with 8 cycles of index reads on an Illumina NovaSeq 6000 platform. Approximately 0.1 million reads per cell were sequenced.

### Data analysis for single-unit RNA-sequencing

Raw sequencing data were aligned to the mouse genome (mm10) and the R package Seurat was used for gene and filtration, normalization, principal component analysis, variable gene finding, clustering analysis, and Uniform Manifold Approximation and Projection (UMAP) dimension reduction. A matrix containing gene-by-unit expression data was imported to create individual Seurat objects. Units with <200 detectable genes and >15% mitochondrial genes were excluded. Data were merged and log-normalized for subsequent analysis. Principal component analysis (PCA) was performed for unbiased clustering. Clusters were visualized with UMAP embedding. Differentially expressed genes between FSC^mid-high^ and FSC^low^ samples in each cluster were determined using the function FindAllMarkers. Gene Ontology (GO) and Kyoto Encyclopedia of Genes and Genomes (KEGG) pathway analyses were performed on differentially expressed genes of each cluster using the gene set enrichment analysis (GSEA) function implemented in the clusterProfiler package; results were plotted using ggplot2.

### Immunostaining of FSC^mid-high^ clusters and colonies

Sorted FSC^mid-high^ clusters or colonies were fixed in 4% paraformaldehyde containing 0.15% Triton X-100 at 4°C overnight, cryoprotected at 4°C overnight in 30% sucrose dissolved in PBS (pH = 7.4-7.6) containing 0.15% Triton X-100 (PBSX), followed by frozen embedding using Optimal Cutting Temperature compound (ThermoFisher). Frozen blocks were sectioned (8 μm thickness) onto glass slides (Fisher Scientific), stored at −80°C, thawed at room temperature (RT), washed with PBSX and blocked with a PBS-based buffer containing 5% donkey serum and 0.1% Triton X-100 for 1 hr at RT. Slides were incubated with primary antibodies at 4°C overnight, washed, treated with secondary antibodies at RT for 2 hrs, washed and finally treated with VECTASHIELD Antifade Mounting Medium following the manufacturer’s instructions (Vector laboratories, Burlingame, CA). Images were captured on a Zeiss Axio-Observer-Z1 microscope (ZEISS) or a Zeiss LSM700. Antibodies used in this study are listed in Supplementary Table 2.

### Histology and staining of pancreas tissue

Pancreata were dissected and fixed in 10% formalin solution for 24 to 72 hrs before being washed with PBSX and stored in 70% ethanol. Samples were paraffin embedded and sectioned (5 μm thickness) onto glass slides from the beginning to the end with representative slides taken 100 μm apart for subsequent quantification. Slides were prepared by baking at 56°C for 3 hrs, followed by de-waxing in xylene bath for 15 min, rehydrated in ethanol, and antigens retrieved using IHC-TekTM Epitope Retrieval Steamer Set (IHCWORLD) for 45 min in sodium citrate buffer (pH=5.5). Slides were then washed with PBSX, treated with permeabilization solution (0.3% Triton X-100 in PBS) for 30 min at room temperature (RT), washed, blocked with a buffer containing 5% donkey serum, 0.1% Triton X-100, and Biogenex Laboratories Power Block (Fisher Scientific) for 2 hrs at RT. Primary antibodies were applied at 4°C overnight and secondary antibodies at RT for 2 hrs. Slides were treated with Vector® TrueVIEW™ Autofluorescence Quenching Kit (Vector laboratories) following manufacturer’s instructions to reduce autofluorescence and unspecific background signals. Nuclei were visualized by incubating for 15 min with 0.1 μg/ml 4,6-diamidino-2-phenylindole (DAPI, Thermo Fisher Scientific) in PBSX. For EdU labeling, Click-iT EdU Alexa Fluor 555 Imaging Kit (Thermo Fisher Scientific) was used following the manufacturer’s instructions. Finally, slides were washed with PBSX and treated with VECTASHIELD Antifade Mounting Medium following the manufacturer’s instructions (Vector laboratories, Burlingame, CA). Images were captured with the ApoTome module on a Zeiss Axio-Observer-Z1 microscope (ZEISS) or a Zeiss LSM880 with Airyscan and processed using Adobe Photoshop and Illustrator 2021. Antibodies used in this study are listed in Supplementary Table 2.

### Transmission electron microscopy (TEM)

Freshly-sorted FSC^mid-high^ clusters were allowed to settle on poly-L-Lysine-treated glass cover slips for 30 min at room temperature. Medium was gently removed and replaced with 0.15 M cacodylate buffer (pH 7.4) containing 2.5% glutaraldehyde and 2 mM calcium chloride as a fixative. After aldehyde fixation, FSC^mid-high^ clusters were rinsed in a 0.15 M sodium cacodylate solution (pH 7.4) containing 2 mM calcium chloride and post-fixed in 1.5% potassium ferrocyanide-reduced 2% osmium tetroxide in 0.15 M cacodylate buffer for 1 hr, rinsed in distilled water and treated with 0.1% aqueous thiocarbohydrazide for 20 min. After further rinsing in distilled water, FSC^mid-high^ clusters were treated with 2% osmium tetroxide for 30 min, rinsed in distilled water, dehydrated in an ethanol series, and infused with Durcupan ACM resin. Ultra-thin sections (~70 nm thick) were cut using a diamond knife on a Leica Ultracut UCT Ultramicrotome (Leica, Wetzlar, Germany) and placed on mesh copper EM grids. Electron microscopy was performed on an FEI Tecnai 12 transmission electron microscope (ThermoFisher) equipped with a Gatan Ultrascan 2K CCD camera (Gatan, Pleasanton, CA).

### Serial block-face 3D scanning electron microscopy (3D-SEM)

Sample blocks were mounted on an aluminum pin and trimmed to 0.5 mm × 0.5 mm. Each specimen was placed in a Zeiss Sigma VP field-emission scanning electron microscope (ZEISS, Oberkochen, Germany) equipped with a serial block-face sectioning unit Gatan 3View (Gatan). A backscatter electron image of the face was obtained under an accelerating voltage of 4 keV and chamber pressure of 20 Pa under the Variable pressure mode. An automatic microtome removed a 70-nm thick slice from the sample and a new image was recorded. This procedure was repeated to yield a data set of 1,000 images from which a complete three-dimensional reconstruction was derived. Images were segmented, analyzed, and visualized using Amira software (ThermoFisher).

### Quantification of DsRed^+^ cells in colonies from Hprt^DsRed/+^ female mice

Images were taken using a Zeiss 700LSM confocal microscope (ZEISS). Samples were imaged using 1-μm optical sections. Optical slices from z-stack-imaged whole colonies were imported into Imaris (Bitplane, Zurich, Switzerland) and backgrounds were corrected. Nuclei diameters were set at 5 μm; red thresholds were set against negative and positive controls. Once parameters were set, a batch analysis was performed on all 32 colonies individually, and DAPI^+^DsRed^+^ or DAPI^+^DsRed^-^ cells were quantified using above annotated batch results.

### Proliferation analysis

Representative pancreatic sections (100 μm apart) were subjected to staining described above and images were taken at 20X magnification with identical channel exposure time for control and injured samples. Images were processed using QuPath v0.2.3 software following the analysis tools described ^64^. Briefly, the object classification function tool was used to quantify single and composite classifier measurements for individual cells (DAPI^+^) that co-expressed Sox9 with Ki67 and/or EdU label. The parameters for object classification (i.e. pixel size, median radius, sigma, minimum area, maximum area, and threshold) were optimized using the Cell Detection algorithm. After optimizing the parameters, a script was generated to perform annotation measurements automatically for all images. Lastly, annotation measurements were exported for analysis (Supplementary Dataset 4).

### Severe Acinar injury mouse model

Tamoxifen (TAM) was injected intraperitoneally into ElaCreERT2;R26^DTR/DTR^ mice at 0.2 mg/g body weight (b.w.) once per day, every other day, for a total of three injections. Three weeks later, diphtheria toxin (DT) (200 ng/ 20g b.w.) was injected once per day for 3 days. For EdU labeling, control and injured adult mice were injected with EdU (100 mg/kg b.w., Abcam, Cambridge, UK) every 24 hrs for 3 days prior to euthanization for tissue collection. Mice were euthanized 3 or 14 days after the last DT injection. Control Cre-;R26^DTR/DTR^ mice received TAM and DT, and control ElaCreERT2;R26 ^DTR/DTR^ mice received TAM and DPBS (vehicle for DT). Pancreata from individual mice were dissected, cleared of fat tissue, dissociated with collagenase B as described above, and total cell number counted. A portion of the dissociated cells/units were stained with anti-CD45 antibodies and analyzed by flow cytometry. Another portion was plated into Matrigel/RSPO1 colony assay; the resulting colonies were counted 3 weeks post-plating.

### Statistical Analysis

Specifics about replicates and statistical tests used in each experiment are available in figure legends or are in the text. For the analysis of two conditions, statistical significance was determined by unpaired, two-tailed Student’s *t*-test. For analysis of more than two conditions, statistical significance was determined by one-way ANOVA, followed by Tukey post hoc analysis. For analysis of conditions with unequal sample size, statistical significance was determined by unpaired, two-tailed Student’s *t*-test with Welch’s correction. Data were statistically analyzed with the GraphPad Prism 8 software and presented as mean ± SD or mean ± SEM. Sample group size (n) is indicated in each figure legend.

## Reporting summary

Further information on research design is available in the Nature Research Reporting Summary linked to this article.

## Data Availability

All underlying data is available to interested researchers within reason.

## Acknowledgments

In memoriam of our beloved colleague Arthur D. Riggs PhD, who passed away during the preparation of this manuscript, we gratefully honor his support and contribution to the current work. We thank Lucy Brown and Alex Spalla for assistance with cell sorting, the integrative genomics core for single-cell RNA-seq analysis, the pathology core for histology analysis, Christiana Crook for technical writing and editing, and Elena C. Chen and Biorender.com for graphic illustration. This work is supported in part by a predoctoral fellowship to J.R.T. from the Norman & Melinda Payson Fellowship at the Irell and Manella Graduate School of Biological Sciences; a predoctoral fellowship to J.O. from the Ford Foundation and from the Helen & Morgan Chu Fellowship at the Irell and Manella Graduate School of Biological Sciences; Juvenile Diabetes Research Foundation postdoctoral fellowship 3PDF2016-174AN and National Pancreas Foundation to J.C.Q.; and National Institutes of Health Grant R01DK099734 to H.T.K. Support from Wanek Family Project of Type 1 Diabetes to H.T.K. is also gratefully acknowledged.

