## Supplementary Figures and Tables for "Identification of a distinct ductal subpopulation with self-renewal and differentiation potential from the adult murine pancreas"

**Supplementary Figure 1:** Gating strategy for fractionation of the parent CD133$^{high}$CD71$^{low}$ population.
**Supplementary Figure 2:** Immunofluorescent control staining for pancreas cell markers.
**Supplementary Figure 3:** Cell-adhesion marker staining of pancreas colonies and controls.
**Supplementary Figure 4:** Quantification of nuclei per individual FSC$^{mid\text{-}high}$ cluster.
**Supplementary Figure 5:** Hprt-dsRed mouse tool validation.
**Supplementary Figure 6:** Drop-Sequencing QC analysis and cluster identification.
**Supplementary Figure 7:** FSC$^{low}$ and FSC$^{mid\text{-}high}$ sub-populations qRT-PCR.
**Supplementary Figure 8:** Top upregulated GO and KEGG pathways among all clusters.
**Supplementary Figure 9:** Pancreas progenitor gene expression pattern among all clusters.
**Supplementary Figure 10:** StemID analysis of Drop-sequencing results and identification of StemID cluster 03 units among all clusters.
**Supplementary Figure 11:** Differentially expressed (DE) genes between cluster 1, TP units, and StemID cluster 03.
**Supplementary Figure 12:** Control samples for acinar injury model and TP cell analysis among normal intercalated, small interlobular, and main ducts.
**Supplementary Figure 13:** Gating strategy for control and injured pancreas samples.
**Supplementary Figure 14:** Male and female cohorts data separated for 3-days after acinar injury.
**Supplementary Figure 15:** Epithelial rosettes are still present 14 days after acinar injury and express ductal makers.
**Supplementary Figure 16:** Graphical abstract.
**Supplementary Table 1:** Taqman probes used for conventional and microfluidic qRT-PCR analyses.
**Supplementary Table 2:** List of antibodies.

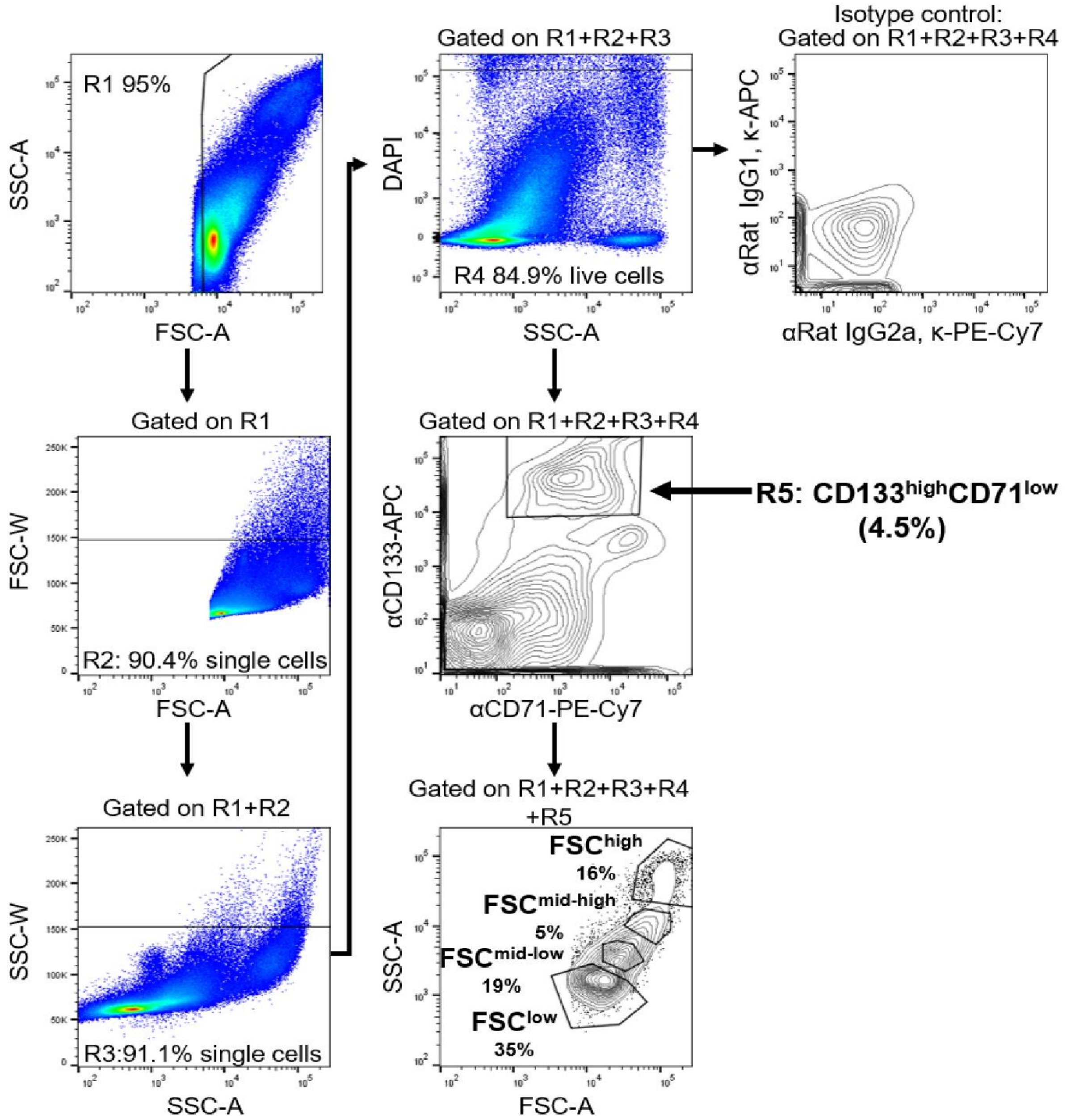

**Supplementary Figure 1. Gating strategy for fractionation of the parent CD133$^{high}$CD71$^{low}$ population.**

Pancreas cells stained for CD133 and CD71 were sequentially gated using the following regions (R): R1 eliminated cell debris; R2 and R3 eliminated cell doublets; R4 eliminated dead cells based on DAPI; R5 gated CD133$^{high}$CD71$^{low}$ cells. R5 cells were further analyzed by size (FSC-A) and granularity (SSC-A), which revealed 4 sub-populations. Percentages of cells in gates are shown from a representative experiment. Abbreviations: FSC-A, forward scatter-area; SSC-A, side scatter-area; FSC-W, forward scatter-width; SSC-W, side scatter-width; APC, Allophycocyanin; PE-Cy7, Phycoerythrin-cyanine dye 7.

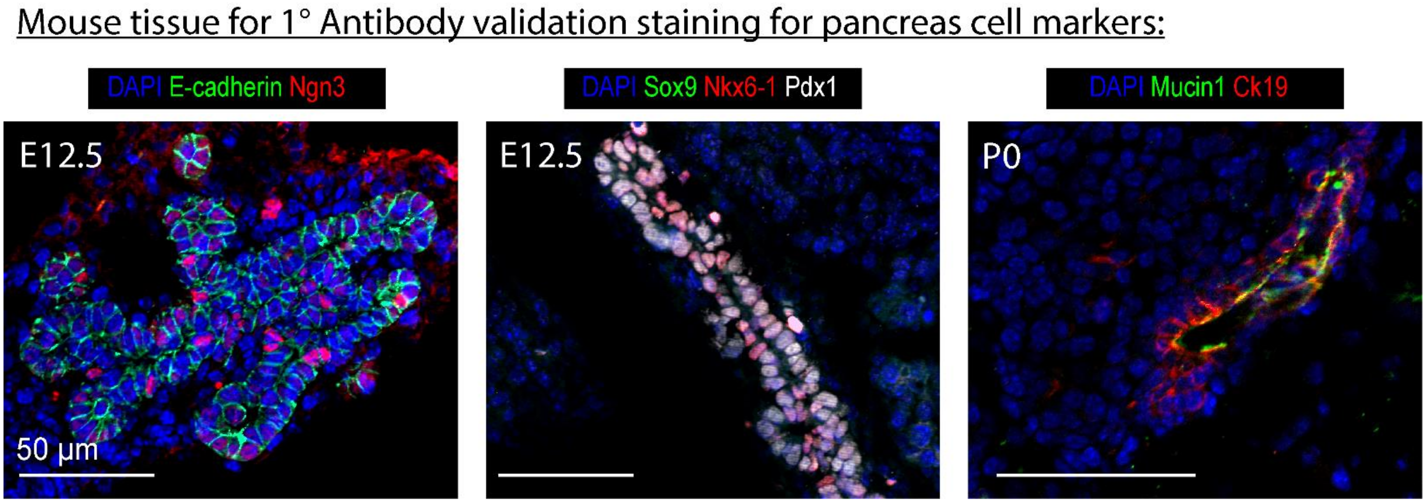

**Supplementary Figure 2. Authentication of antibodies used in immunofluorescence (IF) staining with positive control pancreas tissues**.

Double and triple immunofluorescence (IF) analysis of mouse pancreas at embryonic day 12.5 (E12.5; left and center) and post-natal day 0 (P0; right) were used to validate the expected staining patterns of the primary antibodies that we employed. Scale bars=50 μm.

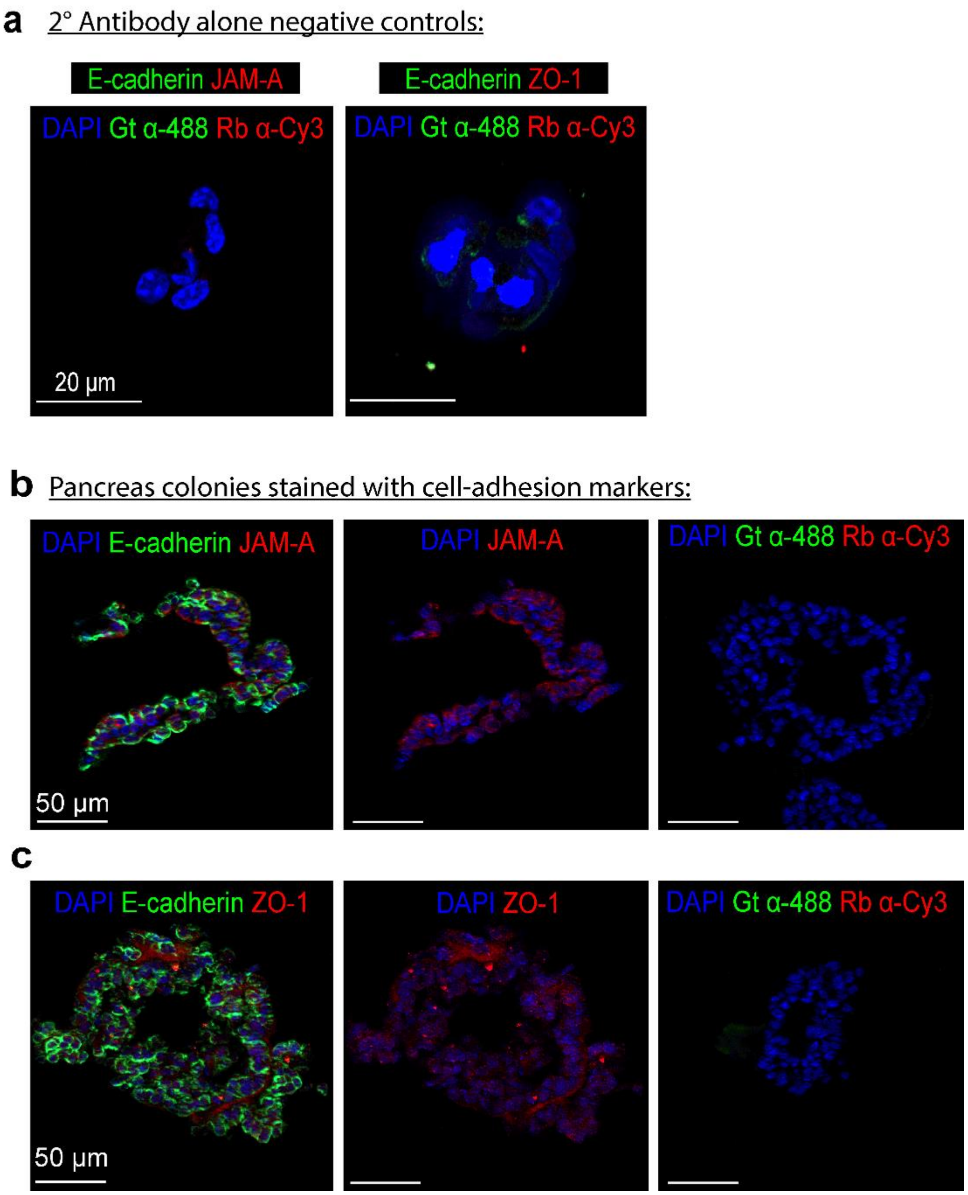

**Supplementary Figure 3. Cell-adhesion marker staining of pancreas colonies and negative control staining.**

(a) Samples treated with secondary antibodies only were used to rule out non-specific staining on $FSC^{mid\text{-}high}$ clusters for E-cadherin (Goat anti-488), JAM-A and ZO-1 (Rabbit anti-Cy3). See also Fig. 4. Scale bars=20 µm. (b-c) IF analysis of $FSC^{mid\text{-}high}$ fraction-derived 3-week-old colonies grown in Matrigel/RSPO1 colony assay. DAPI (blue) identifies individual cells and E-cadherin (green) demarcates cell-to-cell borders. DAPI and E-cadherin were co-stained with either JAM-A (red, b) or ZO-1 (red, c). Negative controls using secondary antibodies only were used to rule out non-specific staining on pancreas colonies for E-cadherin (Goat anti-488), JAM-A and ZO-1 (Rabbit anti-cy3). Scale bars=50 µm.

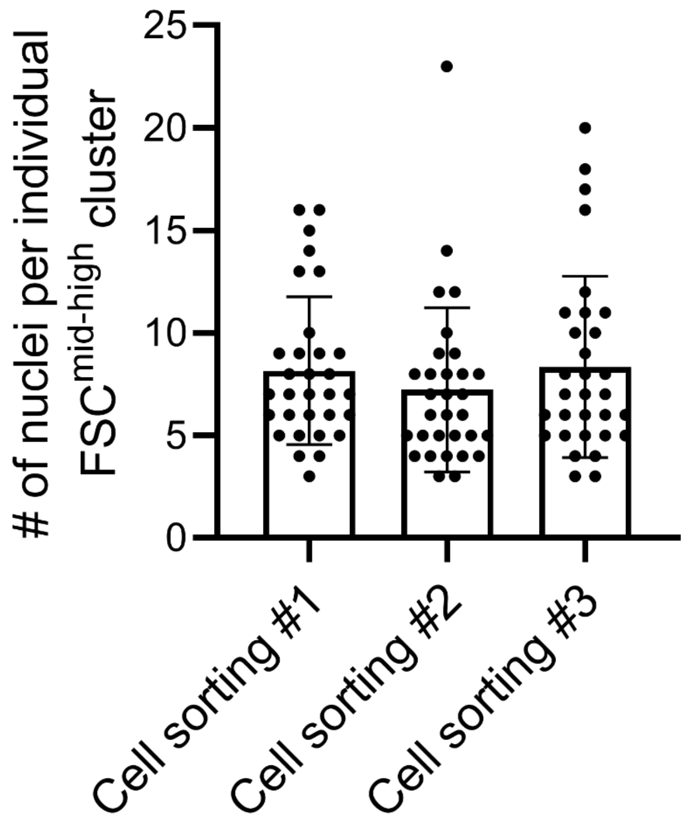

**Supplementary Figure 4. Quantification of nuclei per individual FSC$^{mid-high}$ cluster.**

Three independent sorting experiments were performed to collect FSC$^{mid-high}$ clusters followed by Wright-Giemsa staining to manually quantify the number of nuclei in each cluster. For cell sorting #1 (n=31) the mean number of nuclei was found to be 8.1 ± 3.5 SD. For cell sorting #2 (n=31) the mean number of nuclei was found to be 7.2 ± 3.9 SD. For cell sorting #3 (n=31) the mean number of nuclei was found to be 8.3 ± 4.4 SD. The data from the three independent sorting experiments were combined and shown Fig. 4e. Abbreviation: Standard Deviation (SD).

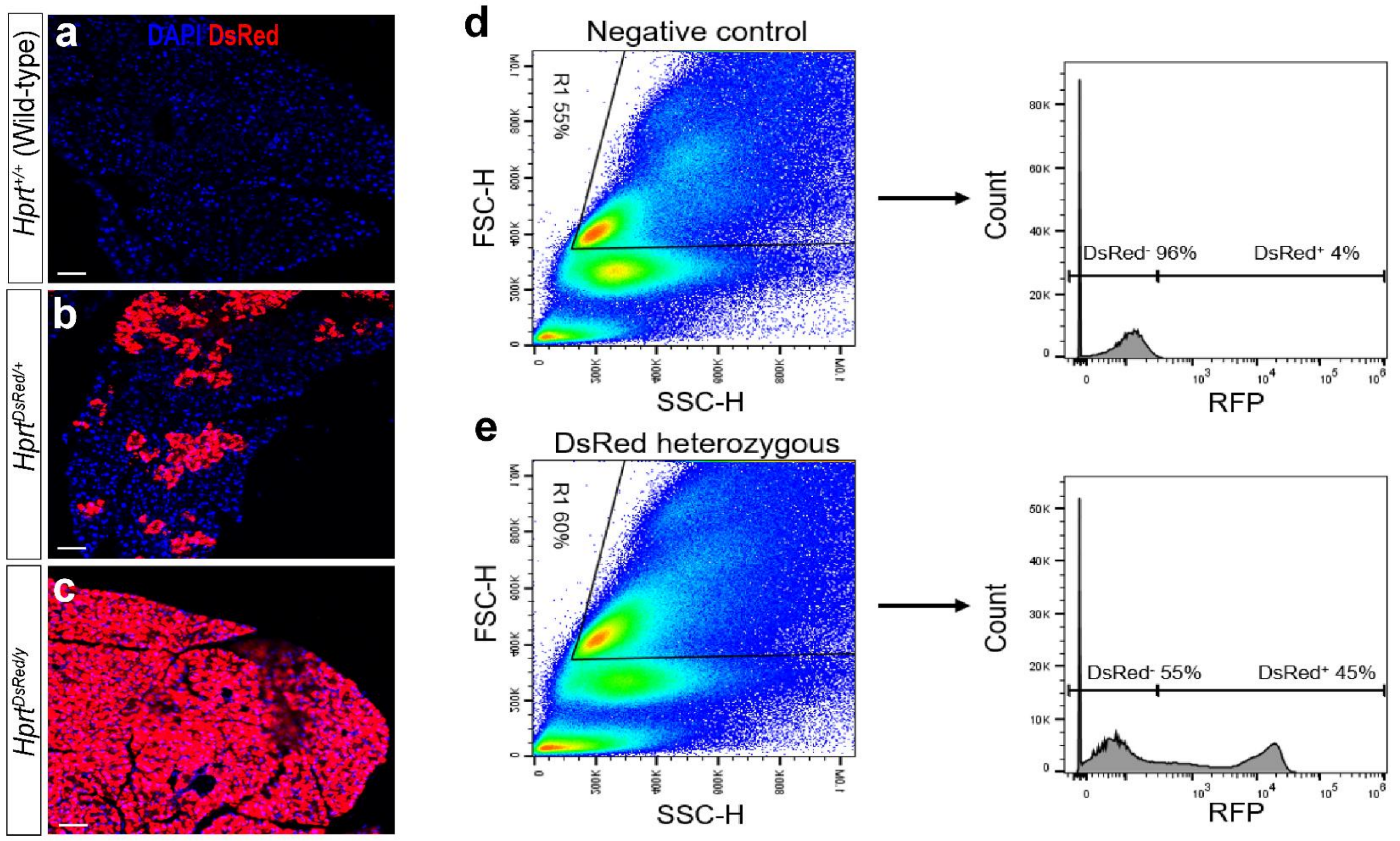

**Supplementary Figure 5. Hprt-dsRed mouse tool validation.**

(a-c) DsRed fluorescence image analysis of pancreata from (a) wild-type female (*Hprt*$^{+/+}$) (negative control, (b) heterozygous female (*Hprt*$^{DsRed/+}$), and (c) homozygous male (*Hprt*$^{DsRed/y}$) mice (positive control). As expected, a mosaic pattern of membrane-bound DsRed labelled-cells (cells in red color) in heterozygous female (*Hprt*$^{DsRed/+}$) was found. (d-e) Spleens were dissected from two 12-week-old female mice and mashed with the plunger end of a syringe to release splenocytes. Splenocytes were filtered, washed, and analyzed using flow cytometry. (d) In wild-type female mice, DsRed staining was present at background levels (4% DsRed$^+$ cells vs. 96% DsRed$^-$ cells). (e) In DsRed heterozygous female mice, approximately half of the splenocytes expressed DsRed (45% DsRed$^+$ cells vs. 55% DsRed$^-$ cells) consistent with a random mosaic pattern.

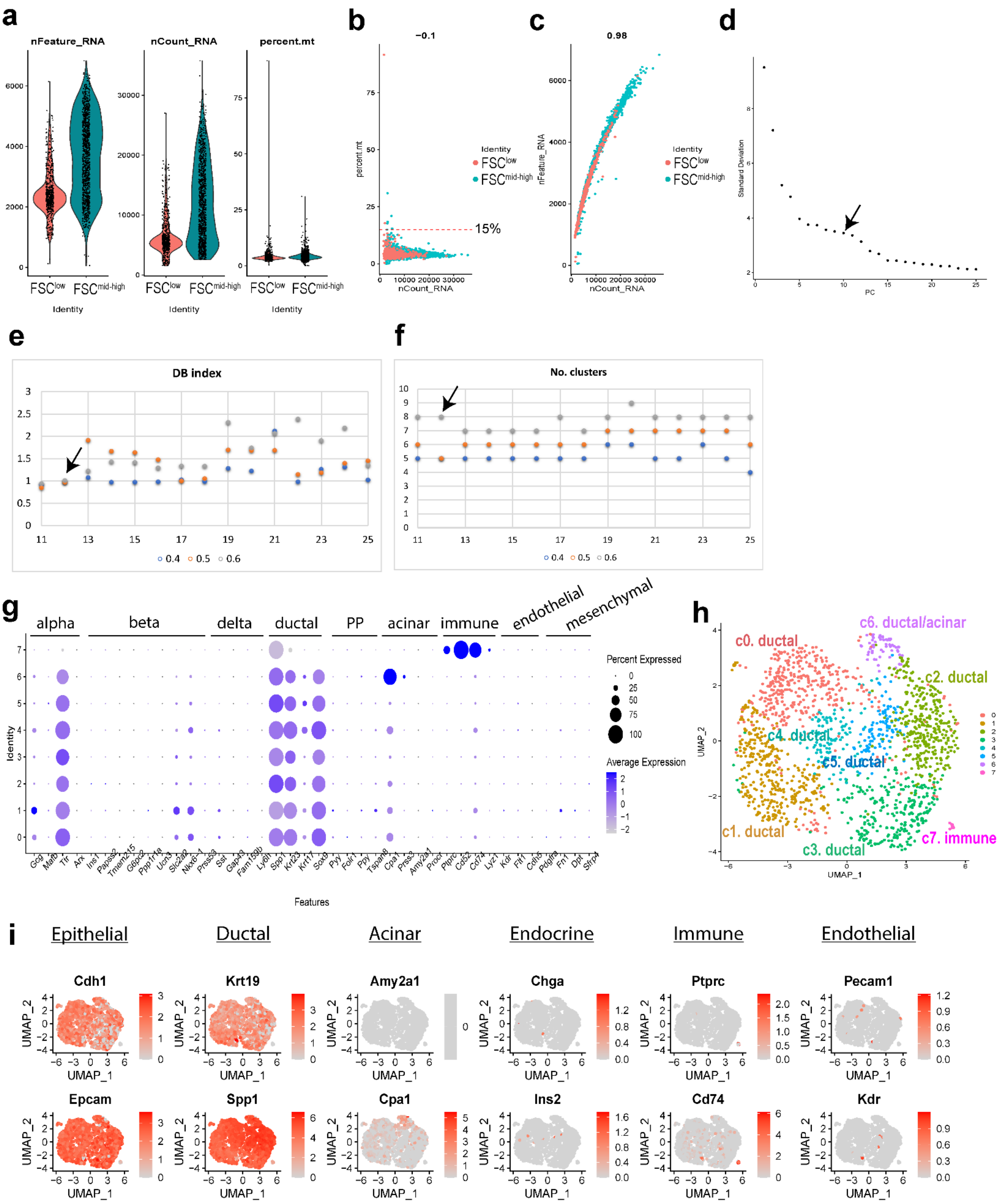

a
nFeature_RNA
nCount_RNA
percent.mt
FSClow
FSCmid-high
Identity
b
-0.1
percent.mt
nCount_RNA
15%
Identity
c
0.98
nFeature_RNA
nCount_RNA
d
Standard Deviation
PC
e
DB index
0.4
0.5
0.6
f
No. clusters
g
alpha
beta
delta
ductal
PP
acinar
immune
endothelial
mesenchymal
Identity
Features
Percent Expressed
Average Expression
Gcg
Mafb
Ttr
Arx
Ins1
Papss2
Tmem215
G6pc2
Ppp1r1a
Ucn3
Slc2a2
Nkx6-1
Prss53
Sst
Gap43
Fam159b
Ly6h
Spp1
Krt23
Krt17
Sox9
Pyy
Folr1
Ppy
Tspan8
Cpa1
Prss3
Amy2a1
Procr
Ptprc
Cd52
Cd74
Lyz1
Kdr
Flt1
Cdh5
Pdgfra
Fn1
Dpt
Sfrp4
h
c0. ductal
c1. ductal
c2. ductal
c3. ductal
c4. ductal
c5. ductal
c6. ductal/acinar
c7. immune
UMAP_1
UMAP_2
i
Epithelial
Ductal
Acinar
Endocrine
Immune
Endothelial
Cdh1
Krt19
Amy2a1
Chga
Ptprc
Pecam1
Epcam
Spp1
Cpa1
Ins2
Cd74
Kdr

**Supplementary Figure 6. Droplet-based RNA-Sequencing quality control (QC) analysis and cluster identification.**

Pancreata from a total of 5 adult (2-4 month old) C57Bl/6 mice were collected, dissociated into a single-cell suspension, stained with antibodies against CD133 and CD71, sorted using a fluorescence-activated cell sorter, and subjected to droplet-based RNA-sequencing using the 10x Genomics platform. Quality control, aggregation, clustering, and analysis was done using Seurat package. (a) The number of feature genes (biologically informative genes) and mRNA counts per unit showed that the sorted $FSC^{low}$ fraction has a unimodal distribution, suggesting that it is a single population. In contrast, the $FSC^{mid\text{-}high}$ fraction appears to have a bimodal distribution, suggesting the presence of both true single cells and clusters. The percentage of mitochondria genes per unit was low in both populations, suggesting a maintenance of plasma membrane integrity after tissue dissociation. (b) Approximately 15% of events displayed higher than 10% mitochondria genes per unit, indicating a compromised plasma membrane. Events that exceeded these criteria were excluded from subsequent analysis. (c) There was a strong positive correlation ($R^2$ = 0.98) between the number of feature genes and the number of mRNAs per unit, suggesting that the mRNAs identified were biologically informative. (d) The elbow plot analysis indicated that 10 principal components were sufficient to capture the majority of the variation in the data. (e) DB index of multiple dimensions indicate that 12 dimensions contain the least variation. (f) Clustering at 12 dimensions with a 0.6x resolution results in 8 clusters (clusters 0 to 7). (g) Bubble plot showing the expression of lineage markers typically found in the pancreas. Ductal lineage genes are found in most of the clusters (0 through 6), with cluster 6 expressing additional acinar marker genes. Cluster 7 expresses immune cell genes. (h) UMAP of clusters 0 to 7 generated with Seurat after combining the datasets for $FSC^{low}$ and $FSC^{mid\text{-}high}$ units. (i) Gene expression levels of various lineage markers in the UMAP plot shown in h.

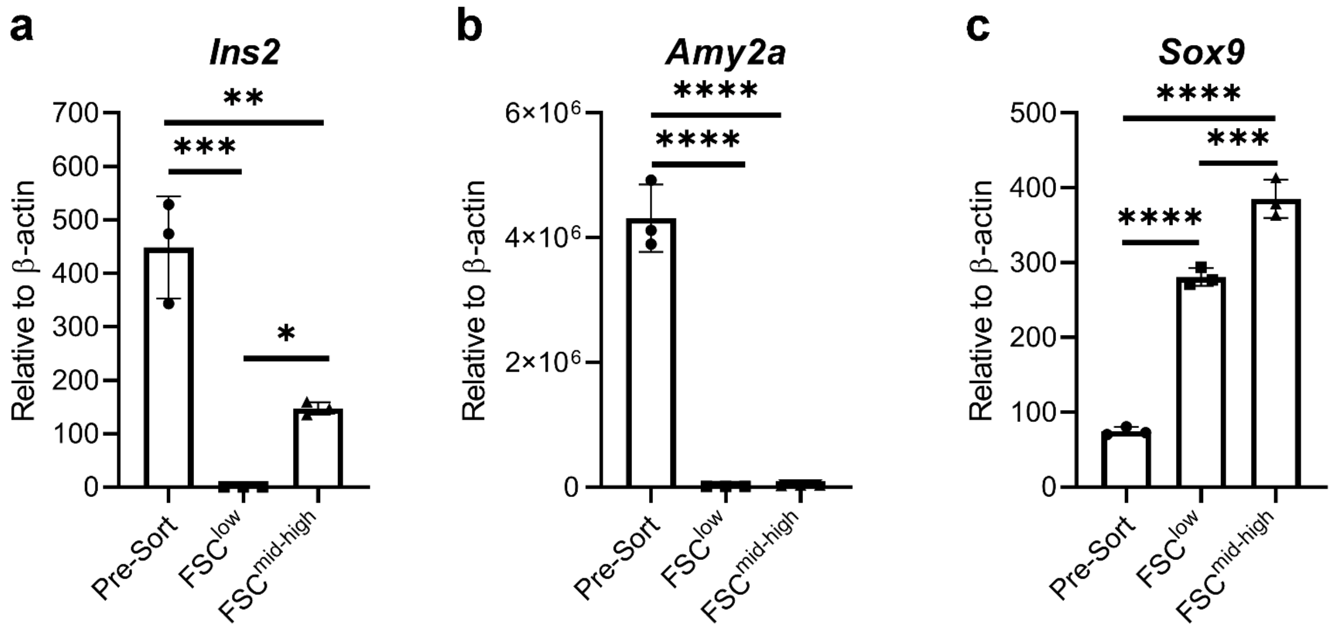

**Supplementary Figure 7. Quantitative RT-PCR analysis of the pre-sorted pancreas cells and after sorting for FSC$^{low}$ and FSC$^{mid-high}$ fractions.**

Pancreata from a total of 5 adult (2-4 month old) C57Bl/6 mice were collected, dissociated into a single-cell suspension, stained with antibodies against CD133 and CD71, FACS sorted, and analyzed for expression of *Insulin 2* (a), *Amylase2a* (b), and *Sox9* (c) using conventional qRT-PCR relative to beta-actin. *$p<0.05$, **$p<0.01$, ***$p<0.001$, ****$p<0.0001$, $n=3$. Statistics were performed using one-way ANOVA multiple comparisons two-tailed Student's t-test.

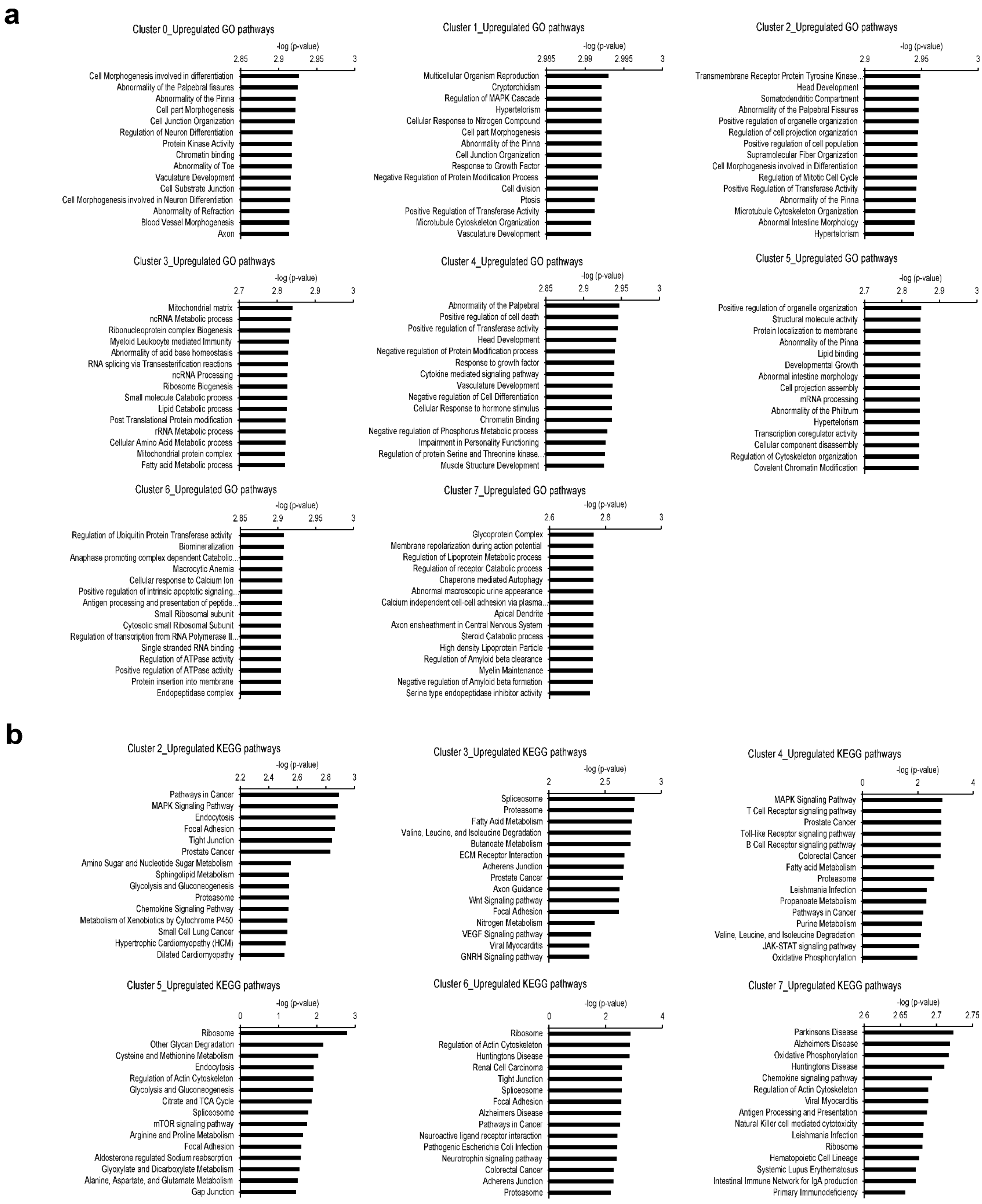
a
Cluster 0_Upregulated GO pathways
-log (p-value)
2.85 2.9 2.95 3
Cell Morphogenesis involved in differentiation
Abnormality of the Palpebral fissures
Abnormality of the Pinna
Cell part Morphogenesis
Cell Junction Organization
Regulation of Neuron Differentiation
Protein Kinase Activity
Chromatin binding
Abnormality of Toe
Vaculature Development
Cell Substrate Junction
Cell Morphogenesis involved in Neuron Differentiation
Abnormality of Refraction
Blood Vessel Morphogenesis
Axon
Cluster 1_Upregulated GO pathways
-log (p-value)
2.985 2.99 2.995 3
Multicellular Organism Reproduction
Cryptorchidism
Regulation of MAPK Cascade
Hypertelorism
Cellular Response to Nitrogen Compound
Cell part Morphogenesis
Abnormality of the Pinna
Cell Junction Organization
Response to Growth Factor
Negative Regulation of Protein Modification Process
Cell division
Ptosis
Positive Regulation of Transferase Activity
Microtubule Cytoskeleton Organization
Vasculature Development
Cluster 2_Upregulated GO pathways
-log (p-value)
2.9 2.95 3
Transmembrane Receptor Protein Tyrosine Kinase...
Head Development
Somatodendritic Compartment
Abnormality of the Palpebral Fissures
Positive regulation of organelle organization
Regulation of cell projection organization
Positive regulation of cell population
Supramolecular Fiber Organization
Cell Morphogenesis involved in Differentiation
Regulation of Mitotic Cell Cycle
Positive Regulation of Transferase Activity
Abnormality of the Pinna
Microtubule Cytoskeleton Organization
Abnormal Intestine Morphology
Hypertelorism
Cluster 3_Upregulated GO pathways
-log (p-value)
2.7 2.8 2.9 3
Mitochondrial matrix
ncRNA Metabolic process
Ribonucleoprotein complex Biogenesis
Myeloid Leukocyte mediated Immunity
Abnormality of acid base homeostasis
RNA splicing via Transesterification reactions
ncRNA Processing
Ribosome Biogenesis
Small molecule Catabolic process
Lipid Catabolic process
Post Translational Protein modification
rRNA Metabolic process
Cellular Amino Acid Metabolic process
Mitochondrial protein complex
Fatty acid Metabolic process
Cluster 4_Upregulated GO pathways
-log (p-value)
2.85 2.9 2.95 3
Abnormality of the Palpebral
Positive regulation of cell death
Positive regulation of Transferase activity
Head Development
Negative regulation of Protein Modification process
Response to growth factor
Cytokine mediated signaling pathway
Vasculature Development
Negative regulation of Cell Differentiation
Cellular Response to hormone stimulus
Chromatin Binding
Negative regulation of Phosphorus Metabolic process
Impairment in Personality Functioning
Regulation of protein Serine and Threonine kinase...
Muscle Structure Development
Cluster 5_Upregulated GO pathways
-log (p-value)
2.7 2.8 2.9 3
Positive regulation of organelle organization
Structural molecule activity
Protein localization to membrane
Abnormality of the Pinna
Lipid binding
Developmental Growth
Abnormal intestine morphology
Cell projection assembly
mRNA processing
Abnormality of the Philtrum
Hypertelorism
Transcription coregulator activity
Cellular component disassembly
Regulation of Cytoskeleton organization
Covalent Chromatin Modification
Cluster 6_Upregulated GO pathways
-log (p-value)
2.85 2.9 2.95 3
Regulation of Ubiquitin Protein Transferase activity
Biomineralization
Anaphase promoting complex dependent Catabolic...
Macrocytic Anemia
Cellular response to Calcium Ion
Positive regulation of intrinsic apoptotic signaling...
Antigen processing and presentation of peptide...
Small Ribosomal subunit
Cytosolic small Ribosomal Subunit
Regulation of transcription from RNA Polymerase II...
Single stranded RNA binding
Regulation of ATPase activity
Positive regulation of ATPase activity
Protein insertion into membrane
Endopeptidase complex
Cluster 7_Upregulated GO pathways
-log (p-value)
2.6 2.8 3
Glycoprotein Complex
Membrane repolarization during action potential
Regulation of Lipoprotein Metabolic process
Regulation of receptor Catabolic process
Chaperone mediated Autophagy
Abnormal macroscopic urine appearance
Calcium independent cell-cell adhesion via plasma...
Apical Dendrite
Axon ensheathment in Central Nervous System
Steroid Catabolic process
High density Lipoprotein Particle
Regulation of Amyloid beta clearance
Myelin Maintenance
Negative regulation of Amyloid beta formation
Serine type endopeptidase inhibitor activity
b
Cluster 2_Upregulated KEGG pathways
-log (p-value)
2.2 2.4 2.6 2.8 3
Pathways in Cancer
MAPK Signaling Pathway
Endocytosis
Focal Adhesion
Tight Junction
Prostate Cancer
Amino Sugar and Nucleotide Sugar Metabolism
Sphingolipid Metabolism
Glycolysis and Gluconeogenesis
Proteasome
Chemokine Signaling Pathway
Metabolism of Xenobiotics by Cytochrome P450
Small Cell Lung Cancer
Hypertrophic Cardiomyopathy (HCM)
Dilated Cardiomyopathy
Cluster 3_Upregulated KEGG pathways
-log (p-value)
2 2.5 3
Spliceosome
Proteasome
Fatty Acid Metabolism
Valine, Leucine, and Isoleucine Degradation
Butanoate Metabolism
ECM Receptor Interaction
Adherens Junction
Prostate Cancer
Axon Guidance
Wnt Signaling pathway
Focal Adhesion
Nitrogen Metabolism
VEGF Signaling pathway
Viral Myocarditis
GNRH Signaling pathway
Cluster 4_Upregulated KEGG pathways
-log (p-value)
0 2 4
MAPK Signaling Pathway
T Cell Receptor signaling pathway
Prostate Cancer
Toll-like Receptor signaling pathway
B Cell Receptor signaling pathway
Colorectal Cancer
Fatty acid Metabolism
Proteasome
Leishmania Infection
Propanoate Metabolism
Pathways in Cancer
Purine Metabolism
Valine, Leucine, and Isoleucine Degradation
JAK-STAT signaling pathway
Oxidative Phosphorylation
Cluster 5_Upregulated KEGG pathways
-log (p-value)
0 1 2 3
Ribosome
Other Glycan Degradation
Cysteine and Methionine Metabolism
Endocytosis
Regulation of Actin Cytoskeleton
Glycolysis and Gluconeogenesis
Citrate and TCA Cycle
Spliceosome
mTOR signaling pathway
Arginine and Proline Metabolism
Focal Adhesion
Aldosterone regulated Sodium reabsorption
Glyoxylate and Dicarboxylate Metabolism
Alanine, Aspartate, and Glutamate Metabolism
Gap Junction
Cluster 6_Upregulated KEGG pathways
-log (p-value)
0 2 4
Ribosome
Regulation of Actin Cytoskeleton
Huntingtons Disease
Renal Cell Carcinoma
Tight Junction
Spliceosome
Focal Adhesion
Alzheimers Disease
Pathways in Cancer
Neuroactive ligand receptor interaction
Pathogenic Escherichia Coli Infection
Neurotrophin signaling pathway
Colorectal Cancer
Adherens Junction
Proteasome
Cluster 7_Upregulated KEGG pathways
-log (p-value)
2.6 2.65 2.7 2.75
Parkinsons Disease
Alzheimers Disease
Oxidative Phosphorylation
Huntingtons Disease
Chemokine signaling pathway
Regulation of Actin Cytoskeleton
Viral Myocarditis
Antigen Processing and Presentation
Natural Killer cell mediated cytotoxicity
Leishmania Infection
Ribosome
Hematopoietic Cell Lineage
Systemic Lupus Erythematosus
Intestinal Immune Network for IgA production
Primary Immunodeficiency

**Supplementary Figure 8. Top upregulated GO and KEGG pathways in each cluster.**

(a) Upregulated Gene Ontology (GO) pathways for clusters 0-7. (b) Upregulated Kyoto Encyclopedia of Genes and Genomes (KEGG) pathways for clusters 2-7. See also Fig. 6 and Supplementary Datasets 2-3.

Pancreas progenitor genes expression pattern among all clusters:

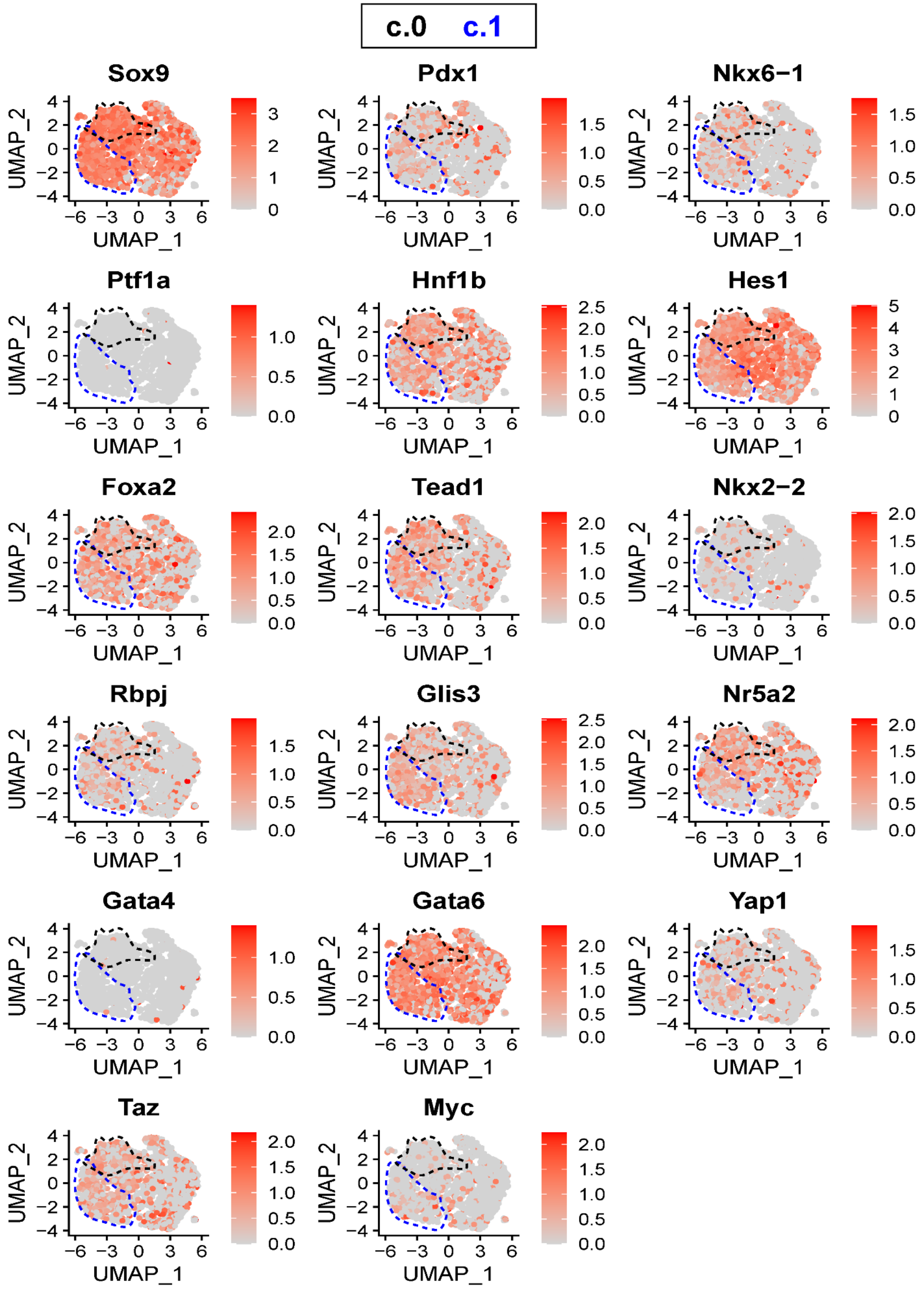

**Supplementary Figure 9: Pancreas progenitor gene expression pattern among all clusters.**

Expression of genes related to progenitor cells during pancreas development are visualized in the UMAPs of the combined datasets from $FSC^{low}$ and $FSC^{mid\text{-}high}$ units with 8 clusters (cluster 0 to 7) identified using Seurat. Clusters 0 (black) and 1 (blue) are outlined in dashed lines. Expression patterns of *Sox9*, *Pdx1*, *Nkx6-1*, *Ptf1a*, *Hnf1b*, *Hes1*, *Foxa2*, *Tead1*, *Nkx2-2*, *Rbpj*, *Glis3*, *Nr5a2*, *Gata4*, *Gata6*, *Yap1*, *Taz*, and *Myc* are shown.

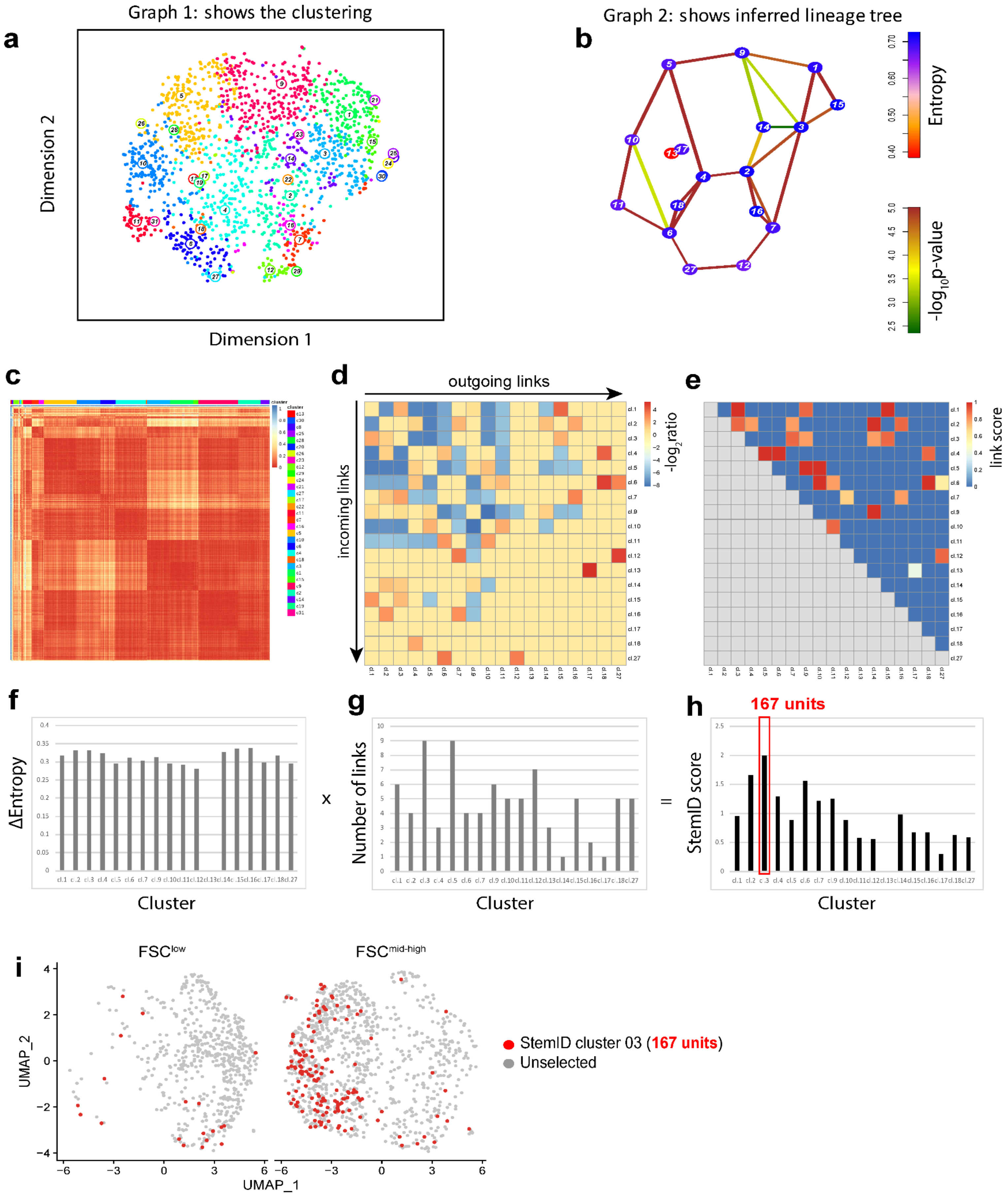

Graph 1: shows the clustering
Graph 2: shows inferred lineage tree
a
b
c
d
e
f
g
h
i
Dimension 2
Dimension 1
Entropy
-log$_{10}$p-value
outgoing links
incoming links
-log$_2$ratio
link score
ΔEntropy
x
Number of links
=
StemID score
167 units
Cluster
FSC$^{low}$
FSC$^{mid-high}$
UMAP_2
UMAP_1
StemID cluster 03 (167 units)
Unselected

**Supplementary Figure 10: StemID analysis of droplet RNA-sequencing datasets and identification of the top StemID group of units among all clusters.**

(a) UMAP of the combined datasets from $FSC^{low}$ and $FSC^{mid-high}$ units that generated 18 clusters using StemID. (b) Graph of the inferred lineage tree. Nodes with multiple lines indicate a higher stemness to the cluster. (c) Heatmap of cell-to-cell transcriptome distances. (d) Heatmap of the incoming vs. outgoing links per cluster. (e) Heatmap of the link score per cluster. (f) Bar graph of the delta entropy (∆Entropy) for each cluster. (g) Bar graph of the number of links for each cluster. (h) Resulting StemID score for each cluster after multiplying ∆Entropy for each cluster with the number of links for each cluster. StemID cluster 03 showed the highest StemID score (1.99) compared to StemID score of the other 17 clusters (1.66 to 0) and therefore the most stem-like population within this dataset. StemID cluster 03 contained 167 units. (i) UMAPs separating $FSC^{low}$ and $FSC^{mid-high}$ units from Seurat analysis. Units from StemID cluster 3 are identified as red dots in the UMAPs from $FSC^{low}$ and $FSC^{mid-high}$ fractions generated using Seurat.

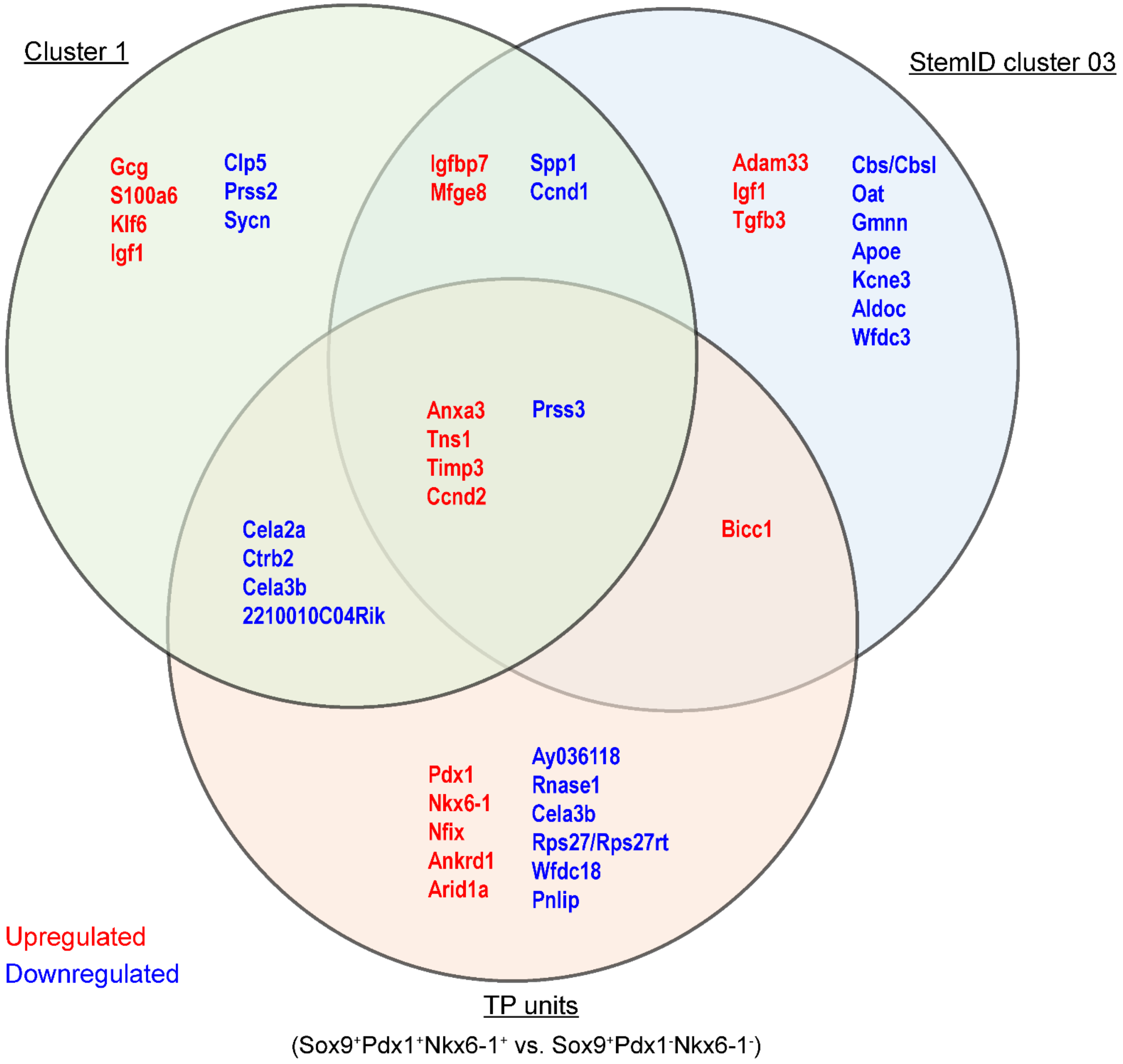

**Supplementary Figure 11: Differentially expressed (DE) genes between Cluster 1, TP units, and StemID cluster 3.**

Venn diagram of the three populations indicating shared upregulated (red) and downregulated (blue) differentially-expressed (DE) genes. See also Supplementary Dataset 1 for complete list of DE genes.

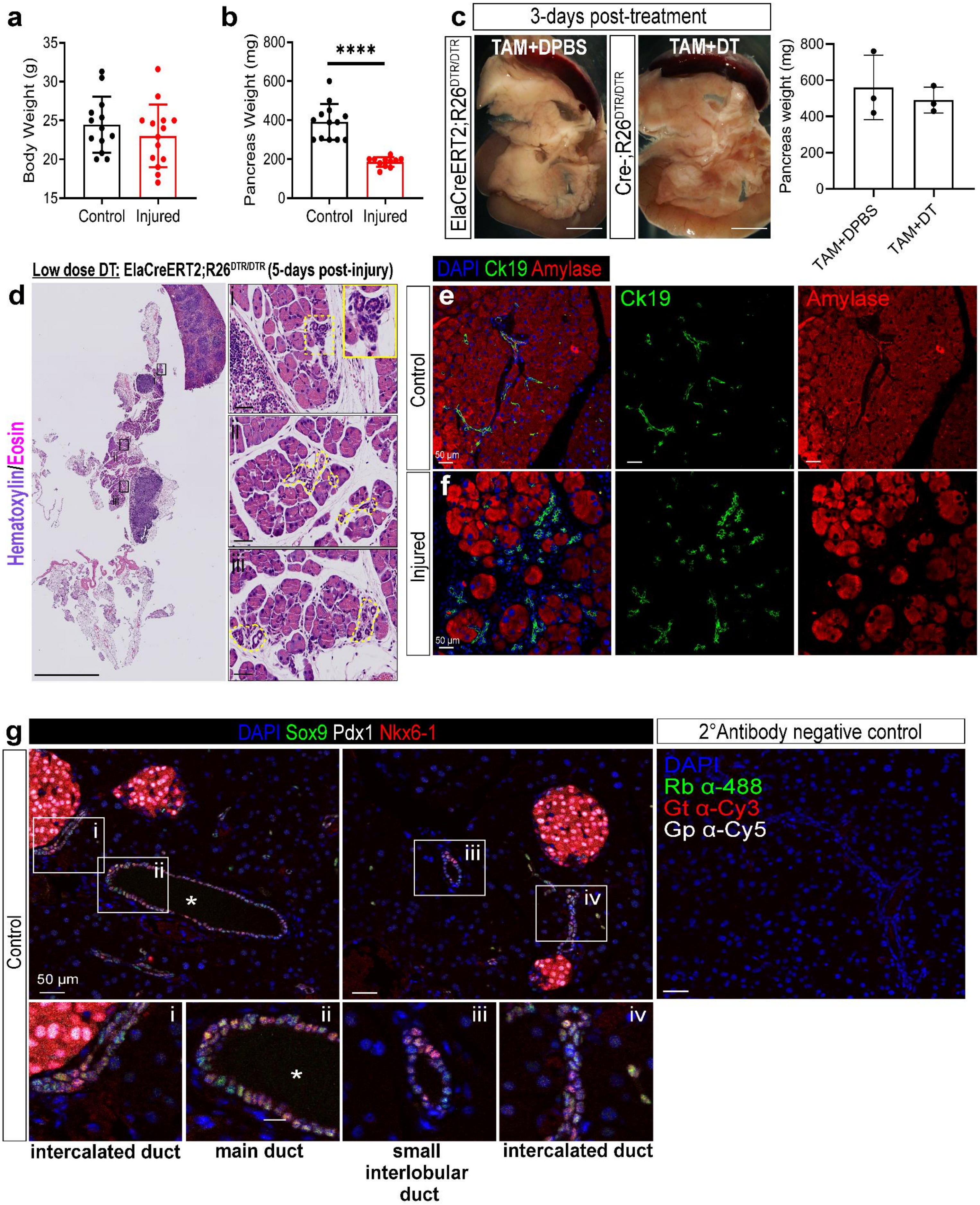
a
Body Weight (g)
Control
Injured
b
Pancreas Weight (mg)
****
Control
Injured
c
3-days post-treatment
ElaCreERT2;R26DTR/DTR
TAM+DPBS
Cre-;R26DTR/DTR
TAM+DT
Pancreas weight (mg)
TAM+DPBS
TAM+DT
Low dose DT: ElaCreERT2;R26DTR/DTR (5-days post-injury)
d
Hematoxylin/Eosin
DAPI Ck19 Amylase
e
Control
Ck19
Amylase
50 μm
f
Injured
50 μm
g
DAPI Sox9 Pdx1 Nkx6-1
2°Antibody negative control
DAPI
Rb α-488
Gt α-Cy3
Gp α-Cy5
Control
50 μm
intercalated duct
main duct
small interlobular duct
intercalated duct

**Supplementary Figure 12: Control samples for acinar injury model and TP cell analysis among normal intercalated, small interlobular, and main ducts.**

(a-c) Brightfield images of dissected pancreas, with spleen and duodenum still attached, from two types of control mice, Cre-positive, top: receiving TAM and DPBS or Cre-negative, bottom: receiving TAM and DT compounds. The pancreas weight of the two types of control mice presented in (c) show no significant difference (right side). n=3. Scale bars=2.5 mm. (a) Body weight of the control and injured mice 3 days after the last DT injection show no significant difference. n=13-14. (b) Pancreas weight (mg) of the control and injured pancreas 3 days after the last DT injection show significant difference. ****$p<0.0001$, n=13-14. (d) H&E staining of the injured pancreas, with a lower dose of DT, 5-days post DT injections. Insets identify ductal rosettes near acinar cells, highlighted in dashed yellow areas. Scale bars=2.5 mm. (e-f) IF analysis of control (e) and injured pancreas (f) co-stained with a duct marker (CK19, green), an acinar marker (Amylase, red), and DAPI (blue). Scale bars=50 µm. (g) IF analysis of normal adult murine pancreas with DAPI (blue), Sox9 (green), Pdx1 (white), and Nkx6-1 (red). Zoomed insets highlight different ducts: main, intercalated, and small interlobular. Negative control (secondary antibodies only) is on the right. Scale bars=50 µm.

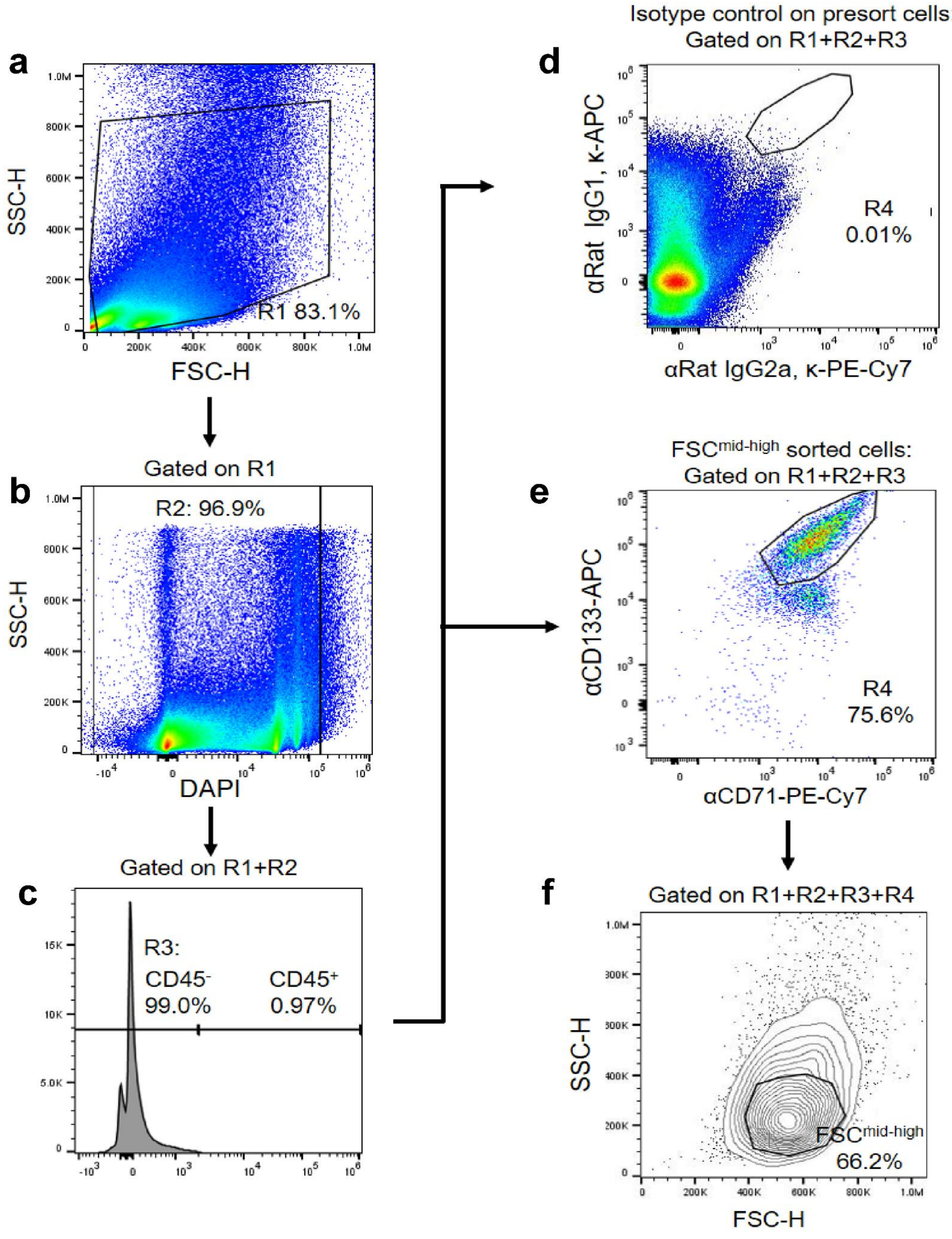

a
R1 83.1%
SSC-H
FSC-H
b
Gated on R1
R2: 96.9%
SSC-H
DAPI
c
Gated on R1+R2
R3:
CD45⁻
99.0%
CD45⁺
0.97%
Isotype control on presort cells:
Gated on R1+R2+R3
d
αRat IgG1, κ-APC
αRat IgG2a, κ-PE-Cy7
R4
0.01%
FSCmid-high sorted cells:
Gated on R1+R2+R3
e
αCD133-APC
αCD71-PE-Cy7
R4
75.6%
f
Gated on R1+R2+R3+R4
SSC-H
FSC-H
FSCmid-high
66.2%

**Supplementary Figure 13: Flow cytometry gating strategy for $CD45^-CD133^{high}CD71^{low}FSC^{mid\text{-}high}$ units.** Gating strategy to identify $CD45^-CD133^{high}CD71^{low}FSC^{mid\text{-}high}$ units on an Attune NxT cytometer using sorted $CD45^-CD133^{high}CD71^{low}FSC^{mid\text{-}high}$ units from an Aria-SORP sorter. Due to differences in hardware between the Attune NxT and Aria-SORP cytometers, sorted $CD45^-CD133^{high}CD71^{low}FSC^{mid\text{-}high}$ units were used as a control to set the $FSC^{mid\text{-}high}$ gate for the Attune NxT cytometer. Briefly, pancreata from 5 wildtype C57Bl/6 mice were dissected, dissociated into a single-cell suspension, and stained with antibodies against CD45, CD133, and CD71. Cells were sorted for $CD45^-CD133^{high}CD71^{low}FSC^{mid\text{-}high}$ units using the Aria-SORP and were analyzed on the Attune NxT cytometer in parallel with unsorted pancreas cells. Cells were sequentially gated using the following regions (R): R1 eliminated cell debris (a); R2 eliminated dead cells (b); R3 eliminated $CD45^+$ cells (c); R4 gated $CD133^{high}CD71^{low}$ cells, showing the isotype (d) and sample (e). $CD133^{high}CD71^{low}$ cells were then analyzed by size (FSC-A) and granularity (SSC-A) to identify the $CD45^-CD133^{high}CD71^{low}FSC^{mid\text{-}high}$ sub-population (f); this gate was used subsequently for analyzing all samples shown in Fig. 7.

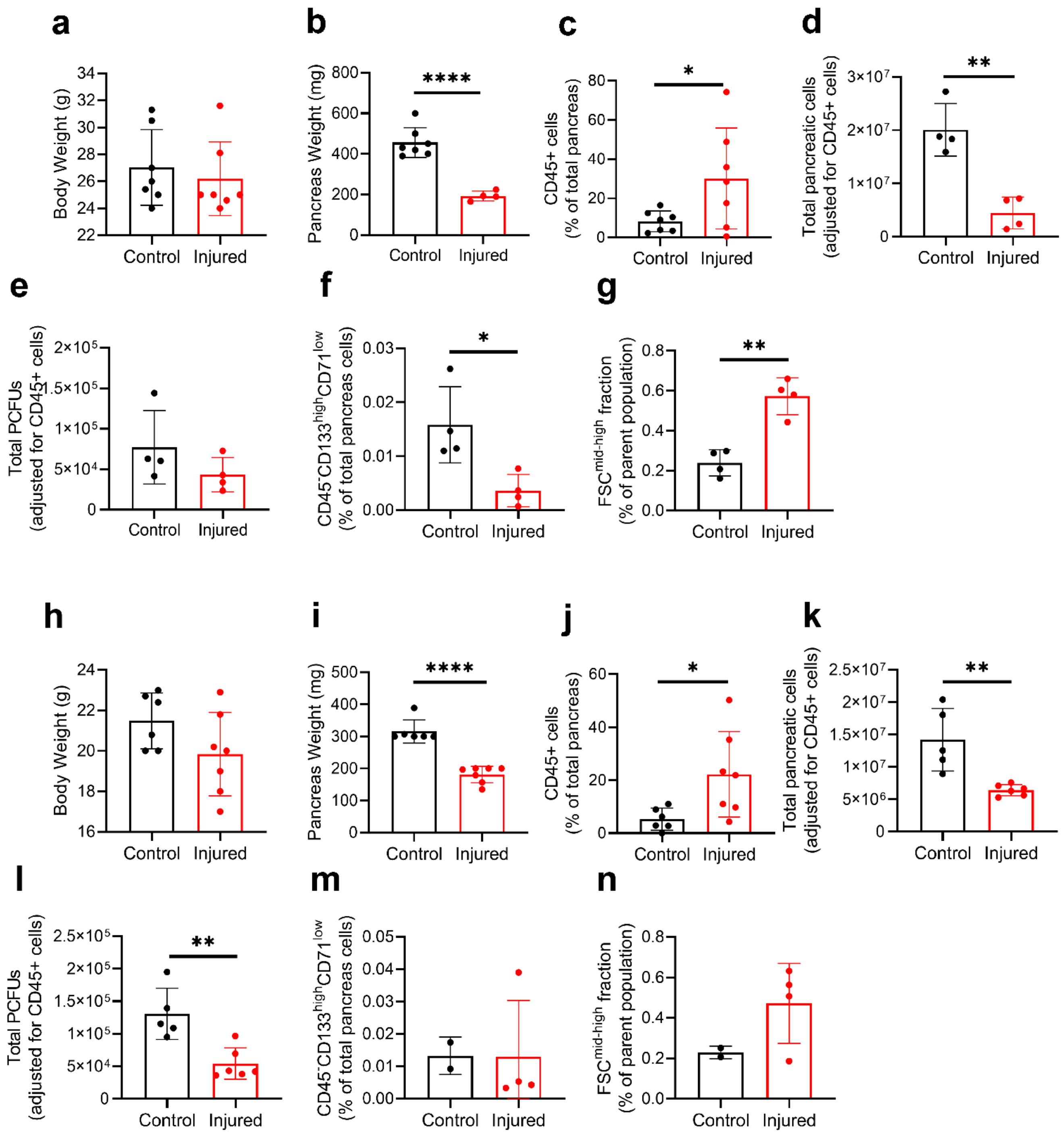

**Supplementary Figure 14: Male and female cohorts data separated for 3-days after acinar injury.**

(a-g) Control and injured male groups. (h-n) Control and injured female groups. *p<0.05, **p<0.01, ****p<0.0001, n=2-7. Statistics were performed using two-tailed Student's t-test Welch's correction. Error bars represent SEM. Male and female cohorts were combined to increase statistical significance, shown in the graphs for Figure 7f-j and Supplementary Fig. 11a-b.

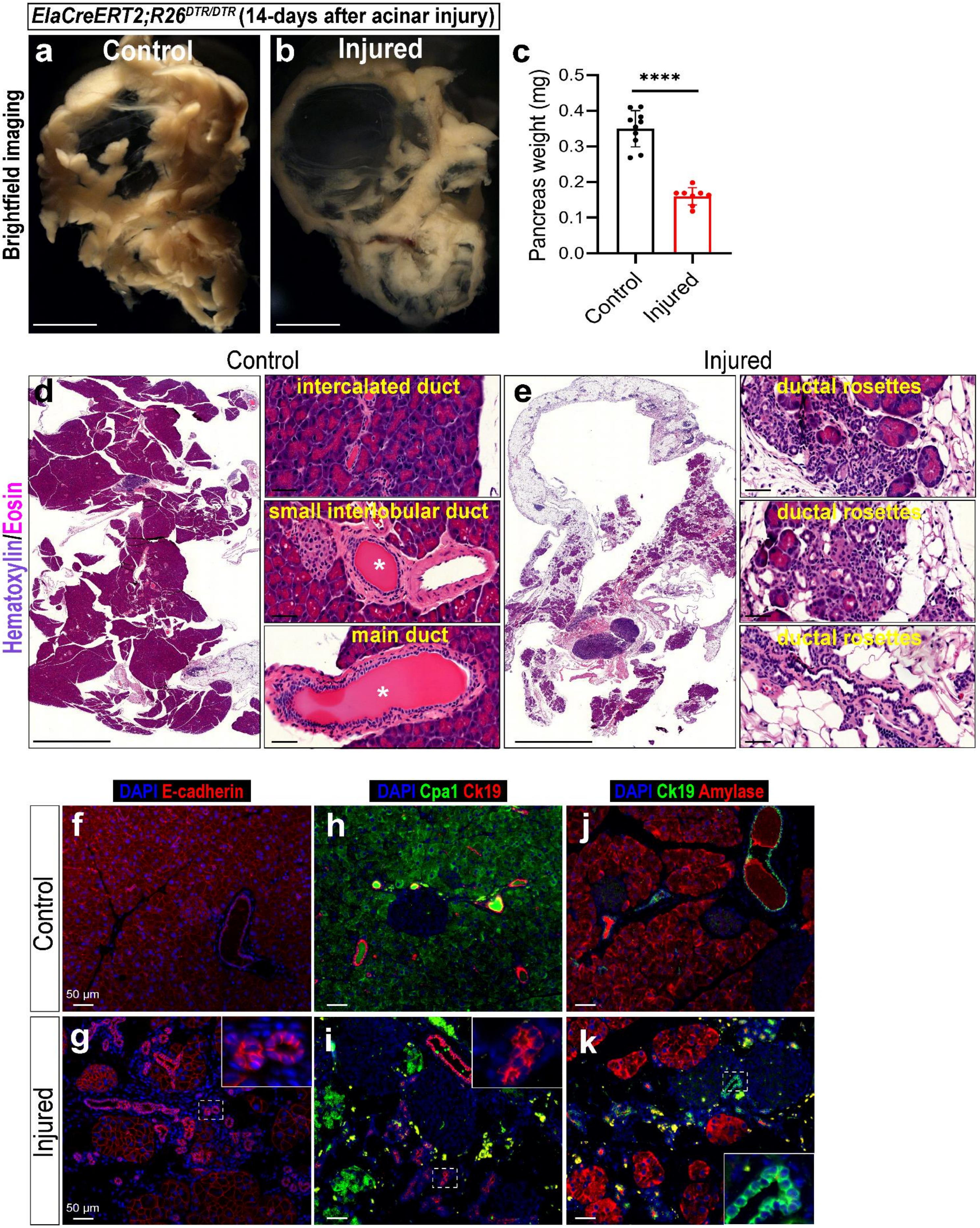

ElaCreERT2;R26DTR/DTR (14-days after acinar injury)
a Control
b Injured
c
Brightfield imaging
Pancreas weight (mg)
0.0
0.1
0.2
0.3
0.4
0.5
****
Control
Injured
Control
Injured
d
e
Hematoxylin/Eosin
intercalated duct
small interlobular duct
main duct
ductal rosettes
ductal rosettes
ductal rosettes
DAPI E-cadherin
DAPI Cpa1 Ck19
DAPI Ck19 Amylase
f
g
h
i
j
k
Control
Injured
50 µm
50 µm

**Supplementary Figure 15: Epithelial rosettes are still present 14-days after acinar injury and express ductal markers.**

(a-b) Brightfield images showing the gross morphology of pancreas samples for control (a) and injured groups (b) 14 days after the last DT injection. (c) Pancreas weight (mg) from control and injured groups show a significant difference 14 days after the last DT injection. ****$p<0.0001$, n=8-10. (d-e) H&E staining of the pancreas from control (d) and injured pancreata (e). Zoomed insets show the normal structure of intercalated, small interlobular, and main ducts in the control pancreas. White asterisk denotes lumen of small interlobular and main ducts. Zoomed insets for injured pancreas show the presence of ductal rosettes. Scale bars=2.5 mm. (f-g) IF analysis of control (f) and injured pancreas (g) co-stained with the epithelial cell marker (E-cadherin, red) and DAPI (blue). (h-i) IF analysis of control (h) and injured pancreas (i) co-stained with markers for duct (CK19, red), acinar (Cpa1, green) and nuclei (DAPI, blue). (j-k) IF analysis of control (j) and injured pancreas (k) co-stained with markers for ductal (CK19, green), acinar (Amylase, red), and nuclei (DAPI, blue). Scale bars=50 µm. Cells in the rosettes of injured pancreas do not simultaneously express duct and acinar markers.

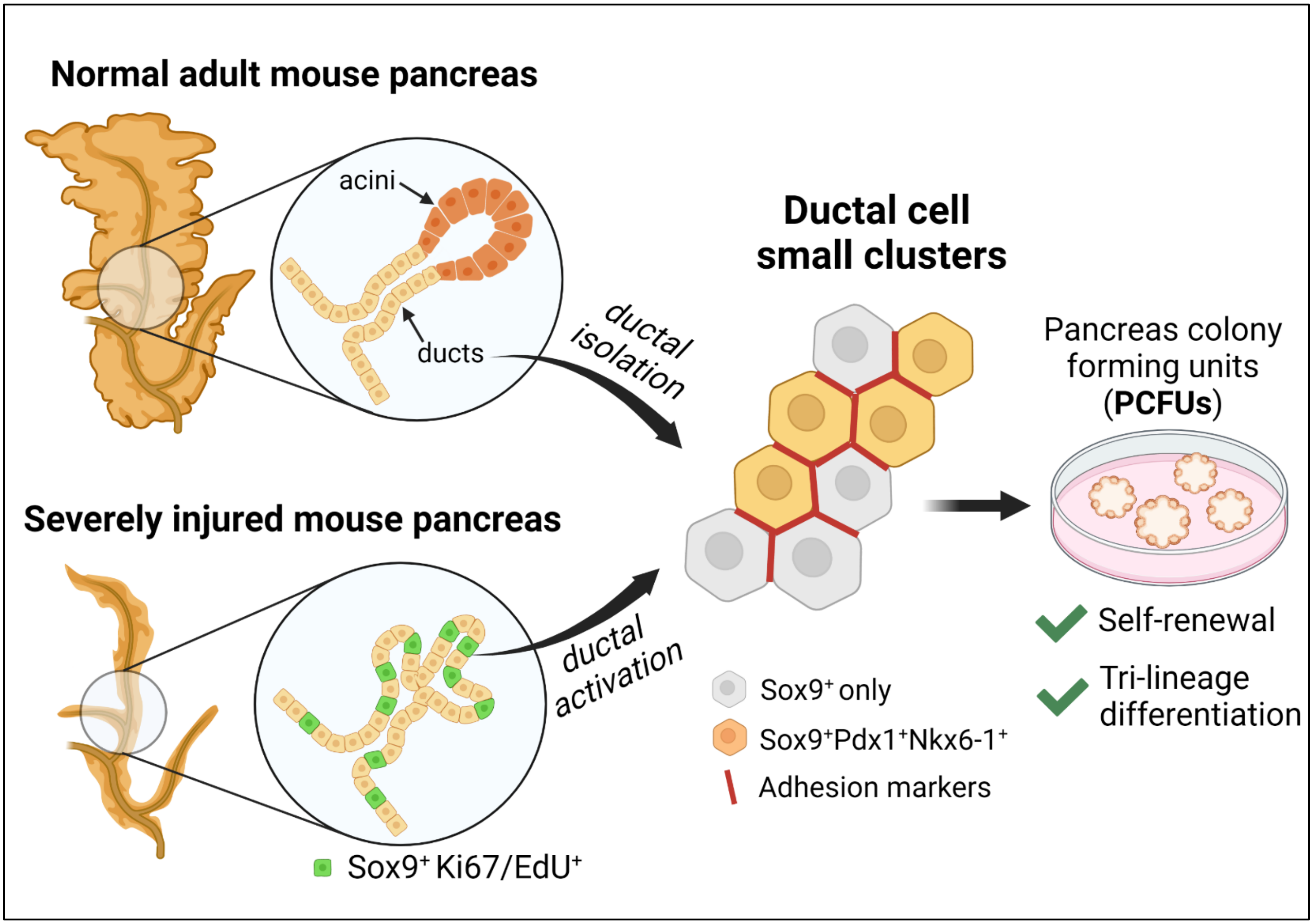

**Supplementary Figure 16: Graphical abstract.**

Adult pancreatic ducts consist of ductal cell clusters (termed FSC$^{mid\text{-}high}$) with self-renwal and differential potential. Ductal cell clusters are enriched for genes involved in cell-cell interactions, organ development, and cancer pathways. Ductal cell clusters exhibit strong cell-adhesion properties that makes them resistant to in vivo acinar injury conditions. Acinar injury conditions induce formation of ductal rosettes that become proliferative within 14 days.

**Supplementary table 1.** Taqman probes used for conventional and microfluidic qRT-PCR analyses.

| **Murine Gene** | **Assay ID from ThermoFisher** |
|---|---|
| *β-actin* | Mm02619580_g1 |
| *Amy2a* | Mm02342487_g1 |
| *Ca2* | Mm00501572_m1 |
| *Prom1* | Mm00477115_m1 |
| *Cpa1* | Mm00465942_m1 |
| *Cela1* | Mm00712898_m1 |
| *Gcg* | Mm00801712_m1 |
| *Hnf6* | Mm00447459_m1 |
| *Ins2* | Mm 00731595_gH |
| *Krt19* | Mm00492980_m1 |
| *Ngn3* | Mm00437606_s1 |
| *Notch2* | Mm00803077_m1 |
| *Pax4* | Mm01159036_m1 |
| *PPY* | Mm01250509_g1 |
| *Ucn3* | Mm00453206_s1 |
| *Slc2a2* | Mm 00446224_m1 |

**Supplementary table 2.** List of antibodies.

| Antibody | Dilution Concentration | | Host | Company | Clone | Cat# | Lot No. |
|---|---|---|---|---|---|---|---|
| | IF | FACS | | | | | |
| C-peptide | 1:100 | | Rabbit | Abcam | Polyclonal | ab1418 | |
| Amylase | 1:300 | | Rabbit | Sigma | Polyclonal | A8273 | |
| CK19 | 1:500 | | Rabbit | Abcam | EP1580Y | ab52625 | |
| CK19 | 1:100 | | Goat | Santa Cruz | Polyclonal | sc-33111 | |
| Neurog3 | 1:100 | | Mouse | DSHB | Monoclonal | F25A1B3 | |
| Mucin 1 | 1:200 | | Armenian Hamster | NeoMarkers | MH1 | HM-1G30-P1 | 1630P1810I |
| CPA-1 | 1:200 | | Goat | R&D Systems | Polyclonal | AF2765 | |
| ZO-1 | 1:300 | | Rabbit | Thermo Fisher | ZMD.437 | 40-2300 | UE289033 |
| JAM-A | 1:300 | | Rabbit | Thermo Fisher | Polyclonal | 36-1700 | UC283843 |
| E-cadherin | 1:300 | | Goat | R&D systems | Polyclonal | AF748 | CYG0518081 |
| Pdx1 | 1:100 | | Guinea pig | Abcam | Polyclonal | ab47308 | GR218270-1 |
| Sox9 | 1:500 | | Rabbit | Abcam | Polyclonal | ab5535 | 3282152 |
| Nkx6-1 | 1:100 | | Mouse | DSHB | Monoclonal | F55A10 | |
| Nkx6-1 | 1:100 | | Goat | R&D Systems | Polyclonal | AF5857 | CDEZ00218121 |
| Ki67-FITC | 1:200 | | Rat | Invitrogen | SolA15 | 11-5698-82 | 2040334 |
| CD133 | 1:100 | | Rat | Millipore Sigma | 13A4 | MAB4310 | |
| CD133-Biotin | | 1:100 | Rat | Invitrogen | 13A4 | 13-1331-82 | 2002734 |
| CD71-PE-Cy7 | | 1:50 | Rat | BioLegend | R17217 | 113812 | B259648 |
| CD45-FITC | | 1:100 | Rat | Thermo Fisher | 30-F11 | 11-0451-82 | 2015766 |
| Streptavidin-APC | | 1:100 | N/A | BioLegend | N/A | 405207 | B288873 |
| Ultra-LEAF purified CD16/32 | | 1:10 | Rat | BioLegend | 93 | 101330 | B287426 |
| Cy™3 AffiniPure F(ab')$_2$ Fragment Donkey Anti-Rabbit IgG (H+L) | 1:2000 (frozen sections)<br>1:500 (paraffin sections) | | Donkey | Jackson ImmunoResearch | | 711-166-152 | |
| Alexa Fluor® 488 AffiniPure Anti-Rabbit IgG (H+L) | 1:1000 (frozen sections)<br>1:500 (paraffin sections) | | Donkey | Jackson ImmunoResearch | | 711-546-152 | |
| Cy™3 AffiniPure Anti-Goat IgG (H+L) | 1:2000 (frozen sections)<br>1:500 (paraffin sections) | | Donkey | Jackson ImmunoResearch | | 705-166-147 | |
| | 1:1000 (frozen sections) | | Donkey | | | 705-545-147 | |

| | | | | | | |
|---|---|---|---|---|---|---|
| Alexa Fluor® 488 AffiniPure Anti-Goat IgG (H+L) | 1:500 (paraffin sections) | | | Jackson ImmunoResearch | | |
| Alexa Fluor® 488 AffiniPure Anti-Guinea Pig IgG (H+L) | 1:1000 (frozen sections)<br>1:500 (paraffin sections) | | Donkey | Jackson ImmunoResearch | | 706-546-148 |
| Alexa Fluor® 647 AffiniPure Anti-Guinea Pig IgG (H+L) | 1:500 (frozen sections)<br>1:500 (paraffin sections) | | Donkey | Jackson ImmunoResearch | | 706-606-148 |
| Cy™3 AffiniPure Anti-Mouse IgG (H+L) | 1:2000 (frozen sections)<br>1:500 (paraffin sections) | | Donkey | Jackson ImmunoResearch | | 715-166-150 |
| Alexa Fluor® 488 AffiniPure Anti-Rat IgG (H+L) | 1:1000 (frozen sections)<br>1:500 (paraffin sections) | | Donkey | Jackson ImmunoResearch | | 712-545-153 |
| Alexa Fluor® 488 AffiniPure Anti-Armenian Hamster IgG (H+L) | 1:1000 (frozen sections)<br>1:500 (paraffin sections) | | Goat | Jackson ImmunoResearch | | 127-545-160 |
| 4,6-diamidino-2-phenylindole (DAPI) | 1:2000 (frozen sections)<br>1:1000 (paraffin sections) | | N/A | Thermo Fisher Scientific | | D1306 |
